# CENP-A/CENP-B uncoupling in the evolutionary reshuffling of centromeres

**DOI:** 10.1101/2024.04.24.590946

**Authors:** Eleonora Cappelletti, Francesca M. Piras, Marialaura Biundo, Elena Raimondi, Solomon G. Nergadze, Elena Giulotto

**Affiliations:** Department of Biology and Biotechnology “Lazzaro Spallanzani”, University of Pavia, Pavia, Italy; Unit of Anatomic Pathology, IRCCS San Matteo Hospital Foundation, Pavia, Italy

**Keywords:** CENP-B, centromere, satellite DNA, karyotype evolution, genus *Equus*

## Abstract

**Background:** While CENP-A is the epigenetic determinant of the centromeric function, the role of CENP-B, the sole centromeric protein binding a specific DNA sequence (CENP-B-box), remains elusive. In the few mammalian species analyzed so far, the CENP-B box is contained in the major satellite repeat that is present at all centromeres. We previously demonstrated that, in the genus *Equus*, some centromeres lack any satellite repeat.

**Results:** Here, we show that, in four *Equus* species, CENP-B is expressed but does not bind the numerous satellite-free and the majority of satellite-based centromeres while it is localized at several ancestral now inactive centromeres. The absence of CENP-B is related to the lack of CENP-B boxes rather than to peculiar features of the protein itself. CENP-B boxes are comprised in a previously undescribed repeat which is not the major satellite bound by CENP-A. Comparative sequence analysis suggests that this satellite was centromeric in the equid ancestor, lost centromeric function during evolution and gave rise to a short CENP-A bound repeat not containing the CENP-B box but being enriched in dyad symmetries. Centromeres lacking CENP-B are functional and recruit normal amounts of the centromeric proteins CENP-A and CENP-C.

**Conclusions:** We propose that the uncoupling between CENP-B and CENP-A may have played a role in the evolutionary reshuffling of equid centromeres. This study provides new insights into the complexity of centromere organization in a largely biodiverse world where the majority of mammalian species still have to be studied.

## BACKGROUND

Centromeres are essential loci required for the correct segregation of chromosomes during cell division. In higher eukaryotes, the DNA component of centromeric chromatin typically consists of tandemly repeated arrays named satellite DNA (1). Despite the well-conserved centromeric function along the evolutionary scale, centromeric satellites are the most rapidly evolving DNA sequences in eukaryotic genomes (2–5). According to the “library hypothesis”, related species share a set of ancestral satellite families that can be differentially modified during the evolution of different lineages (6, 7). New repeats arise and expand in the centromeric core, progressively moving the older units towards the pericentromere, forming layers of different ages (8). During this process, pericentromeric satellites progressively become more and more degenerated and thus cannot be anymore bound by centromeric proteins, avoiding a harmful expansion of the functional centromere (9).

Although satellite DNA is usually associated to centromeres, it is neither sufficient nor necessary for specifying their function (4, 10). Indeed, it is well documented that the centromeric function is not determined by the underlying DNA sequence but rather by the binding of CENP-A, a centromere-specific variant of histone H3, which is the epigenetic marker of functional centromeres (3, 10, 11). CENP-B is the sole centromeric protein that exhibits unequivocal DNA binding specificity (12). The CENP-B target site, called CENP-B box, comprises nine essential nucleotides and represents the only common motif shared by otherwise divergent centromeric satellites (13, 14). The CENP-B protein is highly conserved among mammals and appears to be the result of an exaptation event from a member of the pogo-like transposases family (13–15). The functional domains of CENP-B are the N-terminal DNA-binding region and the C-terminal dimerization domain which are totally conserved in primates and mouse (16, 17). In spite of the conservation of CENP-B and its binding site, several evidences indicate that the protein is dispensable for the centromeric function. Human clinical neocentromeres and Y chromosomes from many species lack CENP-B binding sites, thus they are not bound by CENP-B (2). Conversely, inactive centromeres of pseudo-dicentric chromosomes can retain CENP-B, suggesting that its deposition is not sufficient for centromerization (2). The generation of a human artificial chromosome where CENP-A chromatin was seeded on non-repetitive sequences without the requirement of CENP-B binding (18) confirmed that the absence of CENP-B is compatible with a functional centromere. Moreover, CENP-B knock-out mice are viable, mitotically and meiotically normal demonstrating that CENP-B is not essential for cell division (19–21). However, these animals exhibit low body weight and uterine or testis dysfunctions suggesting a not yet known possible role of CENP-B in the physiology of the reproductive tract (19–21).

The high conservation and dispensability of CENP-B are difficult to reconcile leaving the role of this protein still controversial. It has been proposed that CENP-B might play a role in assembly, disassembly, and/or maintenance of centromere activity (22). The results of several studies suggested that CENP-B may stabilize CENP-A and CENP-C maintenance at centromeres, increasing centromere strength and segregation fidelity of chromosomes (14, 23, 24). Interestingly, loss of the Y chromosome is observed in multiple cancer types, suggesting a high frequency of mis-segregation for this CENP-B negative chromosome (25). It was also proposed that CENP-B participates in the formation of pericentromeric heterochromatin (26) since its depletion causes the disruption of the H3K9me3 environment around centromeres, with subsequent erosion of heterochromatin and genome instability (27, 28). Alternatively, CENP-B conservation might be attributable to non-centromeric functions such as the silencing of transposable elements (29, 30). Finally, it has been proposed that, in centromeric satellites harboring CENP-B boxes, CENP-B mediates the DNA bending required to adopt a non-B conformation typically found at centromeres (31). Recently, it has been proposed that CENP-B may collaborate with CENP-A to establish an open chromatin state by inducing nucleosome DNA unwrapping (32).

In order to shed light on the role of this elusive protein, we investigated CENP-B in the genus *Equus* (horses, asses and zebras). The rapid evolution of these species is marked by exceptionally frequent centromere repositioning events and chromosomal fusions that gave rise to satellite-free centromeres (33–45). In addition, blocks of satellite DNA are often present at non-centromeric chromosome ends, representing relics of ancestral inactivated centromeres or traces of satellite loci exchange (36, 44). The chromosomal distribution of two equid satellite DNA families, 37cen and 2PI, was investigated in horse (*E. caballus*), donkey (*E. asinus*), Grevy’s zebra (*E. grevyi*) and Burchell’s zebra (*E. burchelli*) (36). In the horse (*2n* = 64), all centromeres, with the exception of the one of chromosome 11 (34, 39, 42, 46), are satellite-based and the major centromeric satellite family is 37cen (36, 47). In the donkey (*2n* = 62), 16 satellite-free centromeres are present, while satellite DNA loci are either centromeric or non-centromeric (36, 41). In these two species, satellite-free centromeres derive from repositioning, that is the movement of the centromeric function without DNA sequence modification (48). A high number of chromosomal fusion events led to the karyotypes of the Grevy’s zebra (*2n* = 46) and the Burchell’s zebra (*2n* = 44), where 13 and 15 satellite-free centromeres, respectively, were identified (44). However, in the Grevy’s zebra, the majority of satellite DNA loci are found at non centromeric chromosomal termini, while in the Burchell’s zebra satellite DNA is mainly present at satellite-based centromeres or at fusion sites (36, 44). Thus, the karyotypes of these species represent four different scenarios, providing the opportunity to evaluate the association between CENP-B, centromeres and satellites. Given the coexistence of satellite-free and satellite-based centromeres, the genus *Equus* is an ideal model to study the binding of CENP-B with centromeres and satellite DNA. In this work, we analyzed the binding pattern of CENP-B in these four *Equus* species demonstrating that it is uncoupled from CENP-A binding domains and that the CENP-B box is not contained in the major centromeric satellite. Differently from what previously observed in other systems, in our natural system, the amount of the centromeric proteins CENP-A and CENP-C is not influenced by the presence/absence of CENP-B. These results suggest that the separation of CENP-B from CENP-A may drive the plasticity of equid centromeres.

## RESULTS

### *CENP-B* gene and protein conservation

We retrieved the *CENP-B* gene sequence of the horse (GeneID: 100146426) and the donkey (GeneID: 106847668) from NCBI. The Grevy’s zebra and Burchell’s zebra *CENP-B* genes were identified using BLAT in the genome assemblies Equus_grevyi_HiC and Equus_quagga_HiC, respectively, produced by the DNA Zoo consortium (49). We validated the sequence of the four *CENP-B* genes using both Sanger sequencing and NGS data obtained in our laboratory (Accession Bioproject: PRJNA1054998). Comparative analysis of the DNA sequences and of the deduced protein sequences revealed that CENP-B is highly conserved in the four species with only a few minor differences (Additional file 1: Table S1 and Fig. S1). In particular, the DNA binding and the dimerization domains are identical to the human ones (Additional file 1: Fig. S1), suggesting that the equid CENP-B is functional and able to recognize a canonical CENP-B box.

CENP-B expression was then analyzed in primary fibroblast cell lines from the four species by western blotting using an antibody against the human CENP-B protein. As shown in Figure 1A, the protein is present in all species, although less abundant in Burchell’s zebra, and, in agreement with the intracellular localization of CENP-B in human cell lines (12), resides in the nucleus.

**Figure 1.**
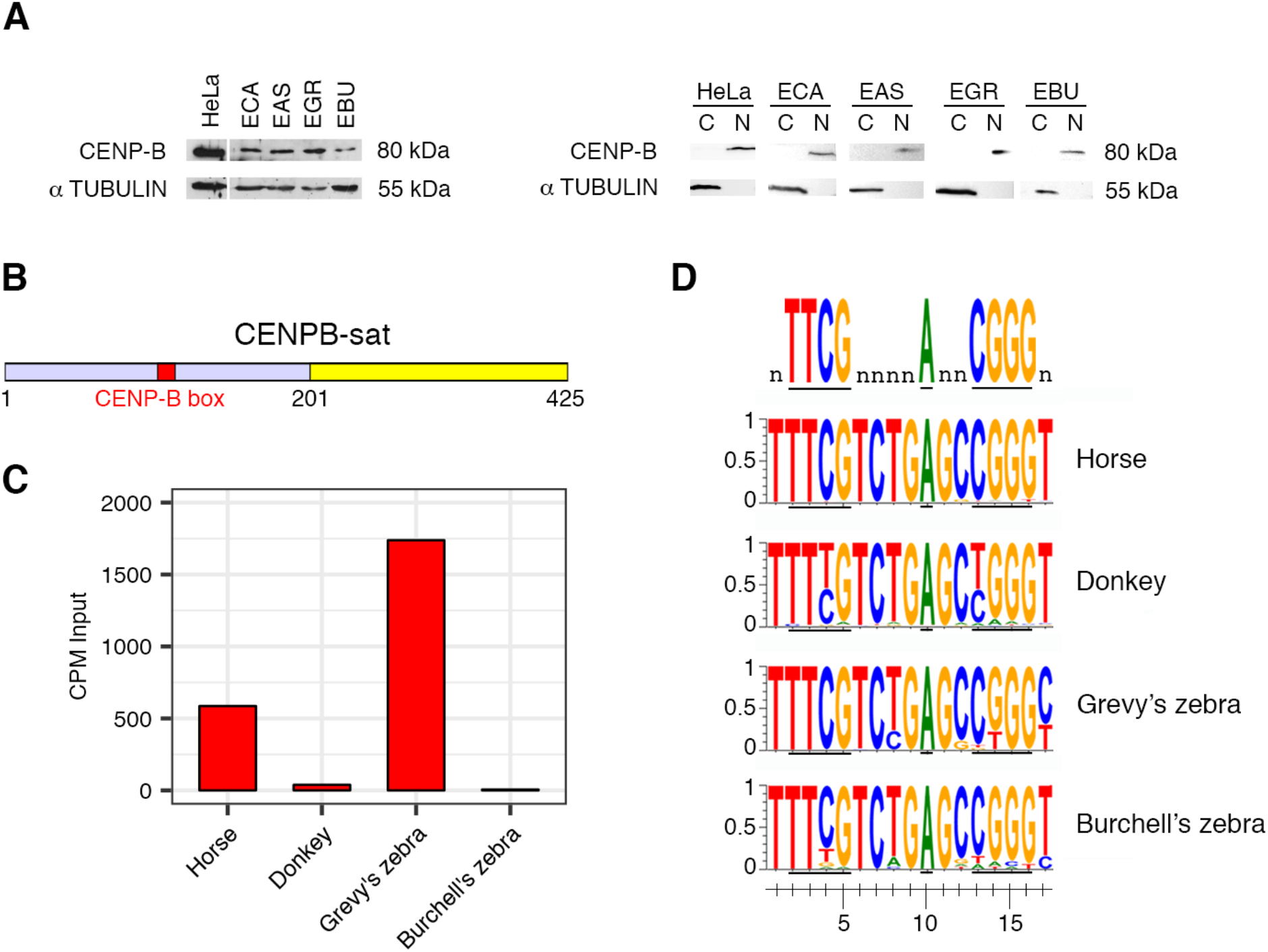
CENP-B protein expression and CENP-B bound satellite in horse, donkey, Grevy’s and Burchell’s zebra. A) Left panel: western blotting on total protein extract from horse (ECA), donkey (EAS), Grevy’s zebra (EGR) and Burchell’s zebra (EBU) with an anti-CENP-B antibody. Protein extracts from human HeLa cells were used as control. All protein extracts were run on the same blot. Right panel: Western blotting on cytoplasmic (C) and nuclear (N) protein extracts from HeLa, horse (ECA), donkey (EAS), Grevy’s zebra (EGR) and Burchell’s zebra (EBU) with an anti-CENP-B 07-735 antibody. Protein extracts of each species were run on different blots. B) Schematic representation of the CENPB-sat satellite sequence. The CENP-B box is colored in red and the region with high identity with the 37cen satellite in yellow. C) Genomic abundance of CENPB-sat in the four species. Values of genomic abundance are reported as Counts Per Million (CPM). D) In the upper row, the 9 nucleotides of the CENP-B box essential for CENP-B binding are shown. The other rows show, for each species, the consensus of the CENP-B box deduced from the Input reads.

### CENP-B binding sites

We previously identified by ChIP-seq, using an anti-CENP-A antibody, one satellite-free centromere in horse (34), 16 in donkey (41), 15 in Burchell’s zebra (44) and 13 in Grevy’s zebra (44). The extraordinarily high number of satellite-free centromeres in equid species raises the question whether CENP-B boxes might be present at such centromeres. We searched for CENP-B boxes (nTTCGnnnnAnnCGGGn) in the genomic sequences of the 45 satellite-free CENP-A binding domains (34, 39, 41, 44) of the four species and did not find any. Therefore, the CENP-B protein is expected not to bind these centromeres.

In all mammalian species analyzed so far, the CENP-B box is comprised within the major centromeric, CENP-A bound, satellite repeat. Surprisingly, the major horse centromeric satellite repeat that we previously identified and characterized, 37cen (47) (SAT_EC in Repbase; AY029358.1 in GenBank) does not contain any CENP-B recognition motif. Moreover, no CENP-B binding sites were detected in the 2PI satellite, the other highly represented satellite DNA family of equid species (36) (ES22 in Repbase), nor in EC137, an accessory pericentromeric satellite DNA element (50) (JX026961.1 in GenBank).

To search for CENP-B binding sites in the horse genome, the best assembled genome sequence among equid species (34, 51), we performed ChIP-seq experiments with an antibody against the human centromeric protein CENP-B on chromatin extracted from horse skin primary fibroblasts. We then aligned the reads with the horse reference genome and identified several peaks in the “unplaced” genomic fraction, which includes highly repetitive DNA sequences lacking chromosomal assignment (Additional file 2: Table S2). These peaks, corresponding to CENP-B binding regions, were contained within arrays of a new satellite family, from now on termed CENPB-sat (Additional file 1: Table S3). CENPB-sat is composed of tandem repeats of a 425 bp unit whose organization is shown in Figure 1B. The GC content of CENPB-sat is 50.5% that is higher than the genomic average (41.0%). Each unit contains a canonical CENP-B box (5’ T<u>TTCG</u>TCTG<u>A</u>GC<u>CGGG</u>T 3’, red in the sketch of Fig. 1B, within a 201 bp fragment unrelated to any other known equine satellite (grey in Fig. 1B). The remaining 224 bps (yellow in Fig. 1B), which do not contain the CENP-B box, share 70% identity with the centromeric satellite 37cen that we previously described (47) (Additional file 1: Fig. S2A). Some CENPB-sat arrays contain degenerated CENP-B boxes or are interrupted by the 22 bp 2PI satellite (Additional file 2: Table S2).

To test the presence of CENP-B box and CENPB-sat in the other equid species, ChIP-seq experiments with the anti-CENP-B antibody were performed on chromatin extracted from skin primary fibroblasts of donkey, Grevy’s zebra and Burchell’s zebra. We evaluated the presence and genomic abundance of CENPB-sat in the four species from the normalized number of reads in the input DNA (Fig. 1C and Additional file 1: Table S4). Grevy’s zebra is the species with the highest genomic representation of CENPB-sat, followed by the horse. In donkey and Burchell’s zebra, CENPB-sat is poorly represented. In Figure 1D, the consensus of the CENP-B box in the four species is shown. These consensus sequences were deduced from the Input reads of each species aligned to the horse CENPB-sat sequence (Additional file 1: Table S3). In the horse, the CENP-B box is highly conserved, indicating that the majority of boxes found in the horse genome are functional. In Grevy’s and Burchell’s zebras, the box is well conserved and only a few mutations were observed in essential nucleotides while, in the donkey, the box is often mutated in two essential nucleotides (C4>T and C13>T).

We then measured the enrichment of CENPB-sat in immunoprecipitated DNA from the four species (Additional file 1: Table S4). As control, we used the ERE-1 retrotransposon, which is well conserved and interspersed throughout the equid genomes and is not expected to be involved in the centromeric function (47, 52). As shown in Additional file 1: Table S4, CENPB-sat is enriched in all immunoprecipitated samples, confirming that, in all species, this satellite is bound by CENP-B. The enrichment of CENPB-sat in Burchell’s zebra indicates that a high fraction of the very few copies of CENPB-sat is bound by CENP-B. The low enrichment of CENPB-sat in the donkey immunoprecipitated chromatin could be due to the fact that only a fraction of the small number of its copies is bound by CENP-B, presumably because of the sequence degeneration (Fig. 1D) that impairs protein recognition. To exclude that the partial identity with 37cen may bias enrichment values, we measured the enrichment of the 201 bp fragment containing the CENP-B box and not sharing any identity with 37cen (Additional file 1: Table S4). Enrichments values of the 201 bp fragment and of the entire CENPB-sat sequence were similar.

We previously showed that 37cen, which does not contain a CENP-B box, is the major CENP-A bound horse centromeric satellite (47). To test whether CENPB-sat is also enriched in CENP-A bound chromatin, we searched for CENPB-sat sequences in ChIP-seq reads that we previously obtained using an anti-CENP-A antibody (41, 44). The results confirmed that, in the horse, 37cen is the most abundant satellite and is enriched in CENP-A chromatin (Additional file 1: Table S5). CENPB-sat is less represented and its enrichment is lower indicating that only a fraction of its few copies is actually bound by CENP-A. In the donkey, 37cen is well represented and bound by CENP-A whereas CENPB-sat is poorly represented, however a high fraction of the very few copies of this satellite reside in centromeric domains. In Grevy’s and Burchell’s zebra, 37cen is nearly absent but its very few copies are contained in CENP-A bound domains. The enrichment of CENPB-sat is very low in the zebras indicating that only a small fraction of this satellite is bound by CENP-A (Additional file 1: Table S5). In other words, in all species, CENPB-sat is not the major centromeric satellite but only a few copies are bound by CENP-A.

A genome wide analysis of the ChIP-seq reads obtained following enrichment with anti-CENP-B antibody from the four species aligned on the horse reference genome allowed us to identify, besides CENPB-sat loci, several enrichment peaks of about 500 bp (Additional file 2: Table S6). These minor peaks, which did not contain any sequence matching satellite repeats, were found in all species, including the horse and Grevy’s zebra where CENPB-sat was well represented and bound by CENP-B. The coordinates of these sites are listed in Additional file 2: Table S7. A subset of these peaks contained one to four CENP-B boxes or CENP-B box-like motifs (at least 7 of the 9 nucleotides essential for CENP-B binding) within single copy sequences. Several peaks mapped in the same position in different species (Additional file 2: Tables S6 and S7). Therefore, CENP-B can bind DNA sequences, not containing CENPB-sat, which are shared among different species. None of these extra-satellite peaks were located within the satellite-free centromeres that we previously described (34, 41, 44).

It is important to remind that CENPB-sat was identified in the horse genome. However, it is well known that satellite sequences are extremely divergent, even among closely related species and that satellite sequences are only partially assembled. To test whether any CENP-B box containing satellite, other than CENPB-sat, is present in the non-caballine species, we retrieved satellite repeats using the TAREAN tool (53) starting from our unassembled reads and did not identify any CENP-B box containing satellite other than CENPB-sat (Additional file 1: Fig. S2B). This search also revealed the presence of the previously identified satellite families (37cen, 2PI, EC137), of two novel satellite repeats (satA, satB) detected in all species, of one repeat shared by donkey and Burchell’s zebra (satC) and of two repeats (satD and satE) that were detected in donkey and Burchell’s zebra, respectively (Additional file 1: Table S8). The consensus sequence of the satellite families identified by TAREAN is reported in Additional file 2: Table S9. We then tested whether any of these satellite repeats were bound by CENP-A taking advantage of our previously published ChIP-seq datasets (41, 44). Using TAREAN and ChIP-seq mapper, we confirmed that, in the horse, the most abundant satellite repeat bound by CENP-A is 37cen (47). In the other species, where the majority of centromeres are satellite-free and satellite DNA is abundant at non-centromeric positions, the organization of the satellites bound by CENP-A is more complex (Additional file 1: Table S8). Some of the novel satellite families, such as satC in donkey and Burchell’s zebra are enriched in immunoprecipitated chromatin.

### Non-canonical DNA structures

It was proposed that non-canonical DNA structures can contribute to centromere specification in the absence of CENP-B binding (31, 54). To test whether this hypothesis may explain the peculiar relationship between CENP-B and centromeric domains in equids, we searched for dyad symmetries and other non-B forming DNA motives in the satellite-free centromeric regions of the four species using the EMBOSS palindrome and nBMST tools. For each species we retrieved the sequence of the CENP-A binding domains (34, 39, 41, 44) and compared their content in non-B structures with those of random genomic regions with the same GC content. As shown in Additional file 1: Fig. S3, we did not detect any enrichment in these sequence features compared to random genomic regions except for A-phased repeats in Grevy’s zebra. In a few cases, we detected lower levels of non-B structures in the centromeric regions. We can conclude that, in the satellite-free centromeres of these species, non-B structures are not relevant. Interestingly, when we performed the same analysis on the consensus sequences of 37cen and CENPB-sat, we observed an enrichment in the number of dyad symmetries in 37cen and in the portion of CENPB-sat sharing high sequence identity with 37cen (Additional file 1: Fig. S4).

### Chromosomal localization of CENP-A, CENP-B and CENP-C proteins and of CENP-B binding satellite

Metaphase spreads from horse, donkey, Grevy’s zebra and Burchell’s zebra were immuno-stained with an anti-CENP-B antibody in two color immunofluorescence experiments with anti-CENP-A or anti-CENP-C antibodies. The localization of the CENP-B binding satellite (CENPB-sat) was then obtained by FISH (Fig. 2). In the four species, all primary constrictions were CENP-A and CENP-C positive, with homogeneous signal intensities, while the distribution of the CENP-B protein and of the CENPB-sat satellite was highly variable and peculiar in each species. The unexpected localization of CENP-B was confirmed using three commercial anti-CENP-B antibodies (Additional file 1: Fig. S5) and different experimental conditions (see Methods).

**Figure 2.**
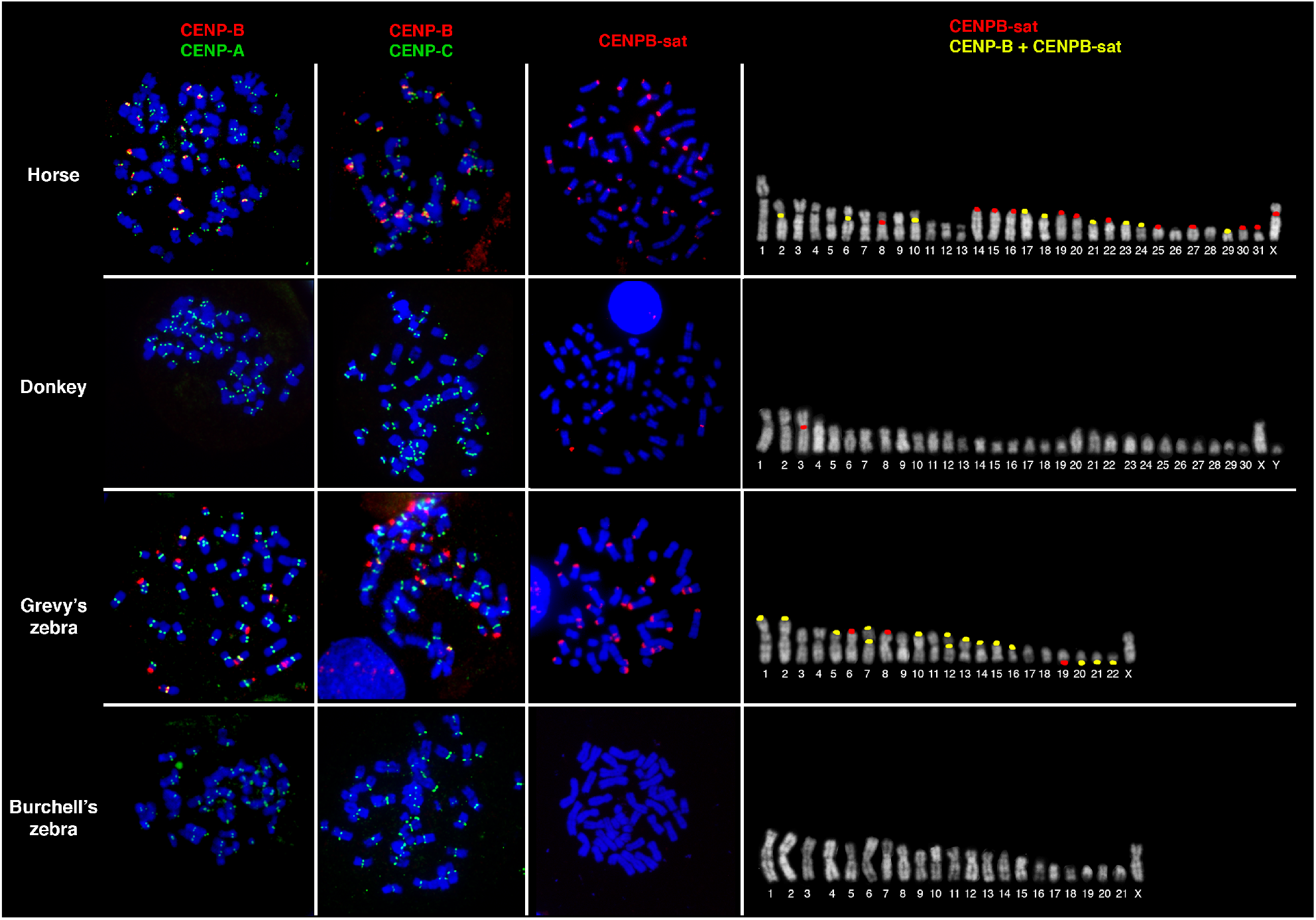
Localization of CENP-B and CENPB-sat in the four species. First column: double immunofluorescence with an anti-CENP-B antibody (red) and an anti-CENP-A serum (green) on DAPI-stained metaphase chromosomes (blue). Second column: double immunofluorescence with an anti-CENP-B antibody (red) and an anti-CENP-C serum (green) on DAPI-stained metaphase chromosomes. Third column: FISH localization of CENPB-sat (red) on DAPI-stained metaphase chromosomes. Fourth column: schematic representation of CENP-B and CENPB-sat signals on metaphase chromosomes. Loci hybridizing with the CENPB-sat probe only are labelled in red. Loci hybridizing with the CENPB-sat probe and positive to CENP-B immunofluorescence are labelled in yellow.

In the horse, the CENP-B protein was detected at the primary constriction of nine out of the 32 chromosome pairs: three metacentric (2, 6 and 10) and six acrocentric chromosomes (17, 18, 21, 23, 24 and 29) (Fig. 2). The signal intensity of CENP-B varied greatly among different chromosomes, and we could not exclude that undetectable amounts of CENP-B might be present also at some additional chromosomes. The CENPB-sat satellite could be detected at the primary constriction of five meta- or submeta-centric chromosomes (2, 6, 8, 10 and X) and sixteen acrocentric chromosomes (14, 15, 16, 17, 18, 19, 20, 21, 22, 23, 24, 25, 27, 29, 30 and 31) (Fig. 2). All CENP-B protein signals colocalized with CENPB-sat signals while, on 12 centromeres, we could detect CENPB-sat signals only (8, 14, 15, 16, 19, 20, 22, 25, 27, 30, 31 and X) (Fig. 2). The lack of detectable CENP-B protein signals at a subset of CENPB-sat positive loci was confirmed by immuno-FISH experiments (Fig. 3A). It is likely that sequence degeneration of the CENP-B box, not detectable by FISH, may prevent binding of the protein at these loci. In addition, we cannot exclude that small amounts of CENP-B protein were present at these loci but were undetectable due to the low resolution of the technique. To confirm the distribution of the CENP-B protein on horse chromosomes, an alternative cellular system, not based on the use of antibodies, was set up. A horse fibroblast cell line, previously immortalized in our laboratory by human telomerase overexpression (55), was transfected with a construct containing the horse *CENP-B* gene tagged with eGFP. As shown in Figure 3B, only a subset of chromosomes was labeled by eGFP signals while most chromosomes lacked detectable eGFP signals. We counted the number of eGFP signals in 10 metaphase spreads. Between fourteen and eighteen signals per metaphase were counted, therefore the distribution of the chimeric protein in transfected cells confirmed the results obtained with the antibodies.

**Figure 3.**
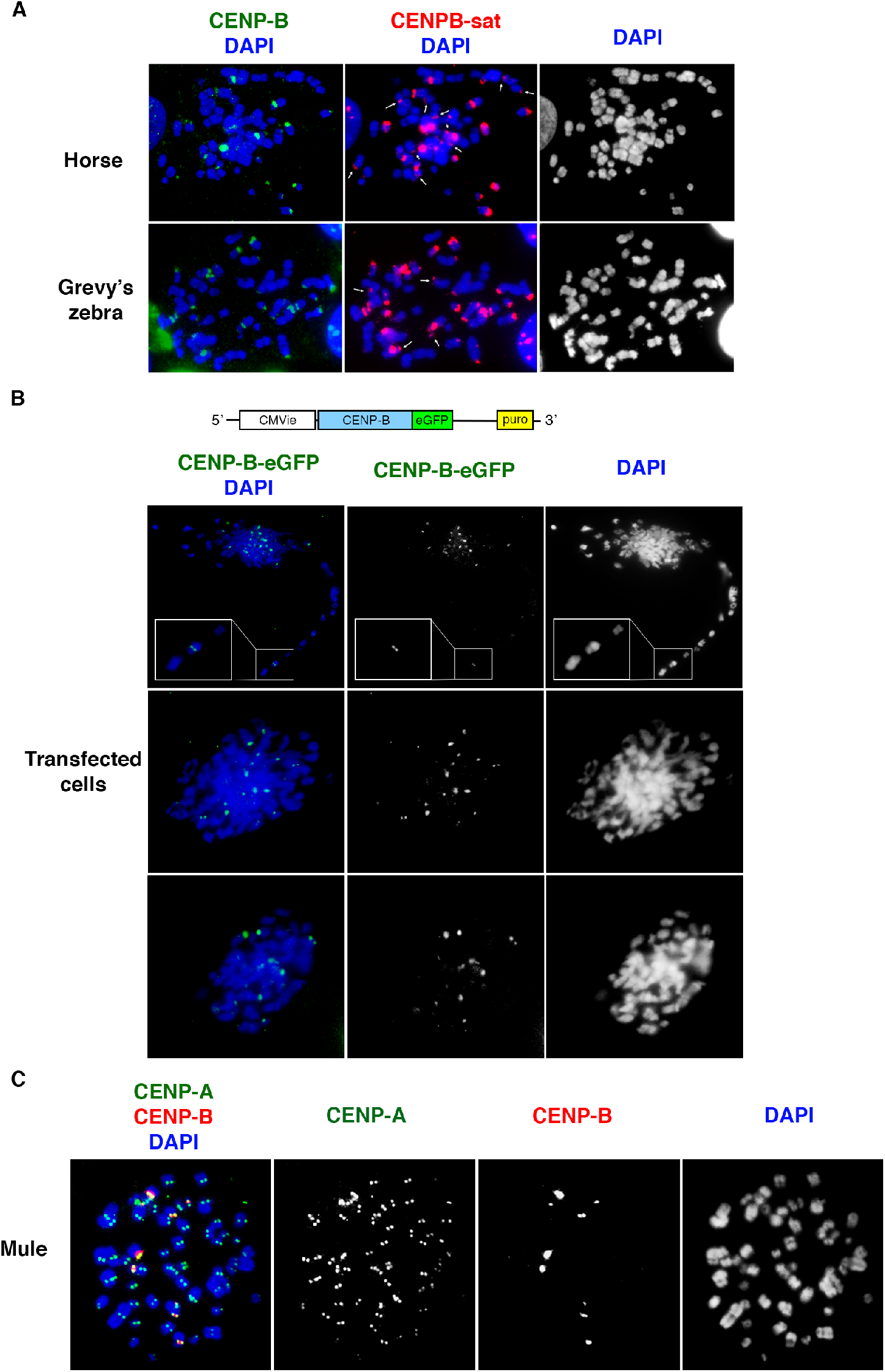
CENP-B binding in horse, Grevy’s zebra and mule. A) Localization of CENP-B protein and CENPB-sat on horse and Grevy’s zebra metaphase chromosomes by immuno-FISH. Left: CENP-B signals (green) on DAPI-stained chromosomes (blue). Middle: FISH CENPB-sat signals (red) on the same metaphase spreads. Immunofluorescence and FISH signals were acquired separately. White arrows point to examples of chromosomes with CENPB-sat but without CENP-B signals. Right: DAPI staining of the same chromosomes. B) A schematic representation of the CENPB-eGFP construct used in transfection is shown on top. Detection of eGFP tagged CENP-B (green) in three horse metaphase spreads. A CENP-B positive and two CENP-B negative chromosomes are boxed and zoomed in the top panel. C) Localization of CENP-A and CENP-B in mule primary fibroblasts. Immunofluorescence with an anti-CENP-A serum (green) and an anti-CENP-B antibody (red) on DAPI-stained metaphase chromosomes.

In the donkey, although all centromeres were labeled by CENP-A and CENP-C, no CENP-B signal could be detected (Fig. 2). This observation indicates that not only the 16 satellite free centromeres but also the 16 satellite-based centromeres are not bound by detectable levels of CENP-B. FISH experiments with the CENPB-sat probe showed hybridization signals on the primary constriction of chromosome 3 only (Fig. 2). However, as in several horse centromeres, the CENP-B protein was not detected on this CENPB-sat positive chromosome. To confirm the lack of CENP-B signal on donkey chromosomes, we examined CENP-B localization in a fibroblast cell line from a mule, that is a hybrid between a horse mare and a donkey jack. Following a double immunofluorescence experiment on metaphase spreads with anti-CENP-B and anti-CENP-A antibodies, specific CENP-B signals were detected only on the primary constriction of the nine chromosomes previously identified in the horse while all other chromosomes, including the complete donkey set, were not labelled (Fig. 3C). Thus, we confirmed that, also in a horse/donkey hybrid cell line, no CENP-B protein signal could be detected on the donkey chromosomes. These results indicate that the absence of CENP-B protein binding at donkey centromeres is related to the lack of CENP-B boxes rather than to peculiar features of the protein itself.

In Grevy’s zebra, CENP-B protein signals were detected at fifteen loci. Surprisingly, only two signals were localized at primary constrictions (chromosomes 7 and 12) while the remaining thirteen were at non-centromeric termini (Fig. 2). In particular, CENP-B localized at a non-centromeric terminus of ten meta- or sub-metacentric (1p, 2p, 5p, 7p, 10p, 12p, 13p, 14p, 15p and 16p) and three acrocentric chromosomes (20q, 21q and 22q). The CENPB-sat and the CENP-B protein colocalized at all sites with the exception of the termini of chromosomes 6, 8 and 19 where only the satellite signal was detected (Fig. 2). The results of immuno-FISH experiments confirm this observation (Fig. 3A). On chromosomes 2, 13, 16 and 19 the terminal non centromeric signals of CENPB-sat and CENP-B showed different intensities on the two homologs suggesting that polymorphism in the copy number of the CENPB-sat repeats may be present in the population (Additional file 1: Fig. S6).

In Burchell’s zebra, as in the donkey, no CENP-B signals could be detected, whereas CENP-A and CENP-C signals were homogeneous on all primary constrictions (Fig. 2). Accordingly, no CENPB-sat hybridization signals were detected in this species, confirming the results of genome sequence analysis (Fig. 1C).

We then performed 3D-immunofluorescence experiments using anti-CENP-B and anti-tubulin antibodies (Fig. 4, Additional file 3-10: movies S1-S8). With this methodology cell morphology is preserved as opposed to the method used to prepare metaphase spreads (Fig. 2). Since it is well known that CENP-B localizes at all human centromeres, HeLa cells were used as control. As expected, in HeLa cells, CENP-B fluorescence was present in the nucleus, with discrete foci corresponding to centromeres (Fig. 4A, Additional file 3: movie S1). In horse and Grevy’s zebra the situation was similar, with discrete nuclear CENP-B foci (Fig. 4A, Additional files 4-6: movies S2-S4). On the contrary, donkey and Burchell’s zebra lack CENP-B nuclear foci and only a diffuse fluorescence was observed (Fig. 4A, Additional files 7-10: movies S5-S8). This result is consistent with the absence of chromosomal CENP-B loci detectable by immunofluorescence (Fig. 2). The presence of discrete chromosomal CENP-B loci only in horse and Grevy’s zebra was also observed in metaphase cells (Fig. 4B, Additional files 5-6: movie S3 and S4) confirming the results obtained with metaphase spreads (Fig. 2).

**Figure. 4.**
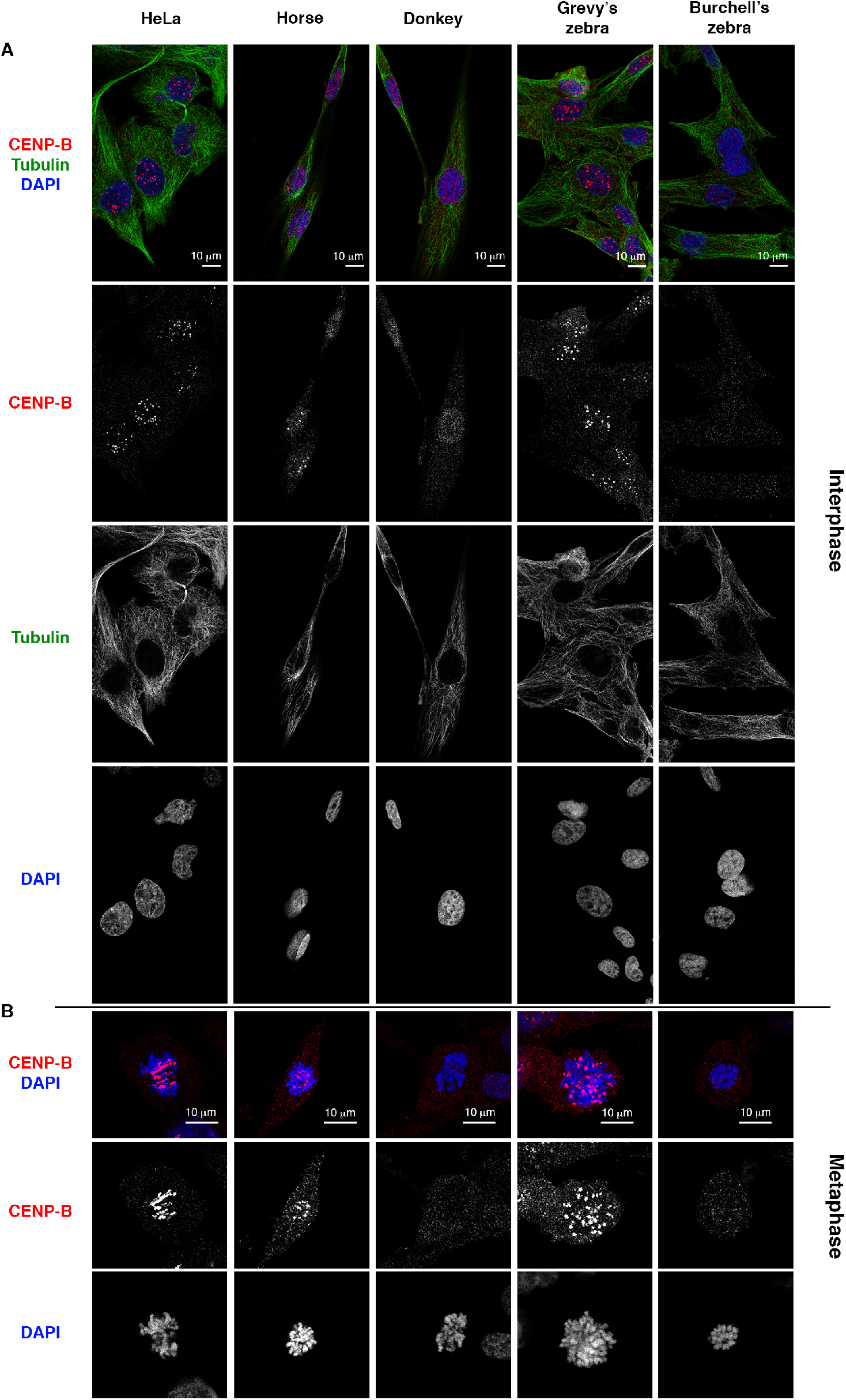
Localization of CENP-B in interphase and metaphase cells by 3D immunofluorescence. A) Optical sections from 3D-immunofluorescence with anti-CENP-B (red) and an anti-tubulin (green) antibodies on whole cells in HeLa, horse, donkey, Grevy’s zebra and Burchell’s zebra. B) Optical sections from 3D-immunofluorescence with an anti-CENP-B antibody (red) on metaphase cells in HeLa, horse, donkey, Grevy’s zebra and Burchell’s zebra. Nuclei were counterstained with DAPI (blue). Bars = 10 μm.

### CENP-B positive and negative chromosomes: CENP-A and CENP-C quantification and segregation fidelity

It has been proposed that the amount of CENP-B protein at centromeres directly correlates with the amount of CENP-C, resulting in different degrees of centromere stability (14).

Taking advantage of the presence of CENP-B positive and CENP-B negative centromeres in the horse, we tested the possible correlation among the levels of the three centromeric proteins by immunofluorescence. As shown in Figure 5A and B, the centromeric CENP-B signals did not show the typical speckled pattern of CENP-A and CENP-C but were broad extending over the pericentromeric area, confirming that CENP-B is localized outside the centromeric core. In addition, this Figure clearly shows that the intensity of CENP-A and CENP-C signals is homogeneous regardless the presence or absence of CENP-B signals. We then measured fluorescence intensities of CENP-A, CENP-B and CENP-C signals using the program Fiji. As shown in Figure 5C, CENP-A and CENP-C signals did not differ in CENP-B positive and negative centromeres according to Student’s t test (t-value = 1.47851 and p-value = 0.140208 for CENP-A; t-value = 0.8983 and p-value = 0.185017 for CENP-C). Moreover, according to Spearman’s correlation test, fluorescence intensity of CENP-B signals was not correlated with signal intensity of CENP-A (R = 0.0675 and p-value = 0.215856) and CENP-C (R = 0.0536 and p-value = 0.432112 for CENP-C). Thus, differently from previous results in human and mouse (14), the levels of CENP-A and CENP-C were rather homogeneous and independent from the presence and amount of CENP-B.

**Figure 5.**
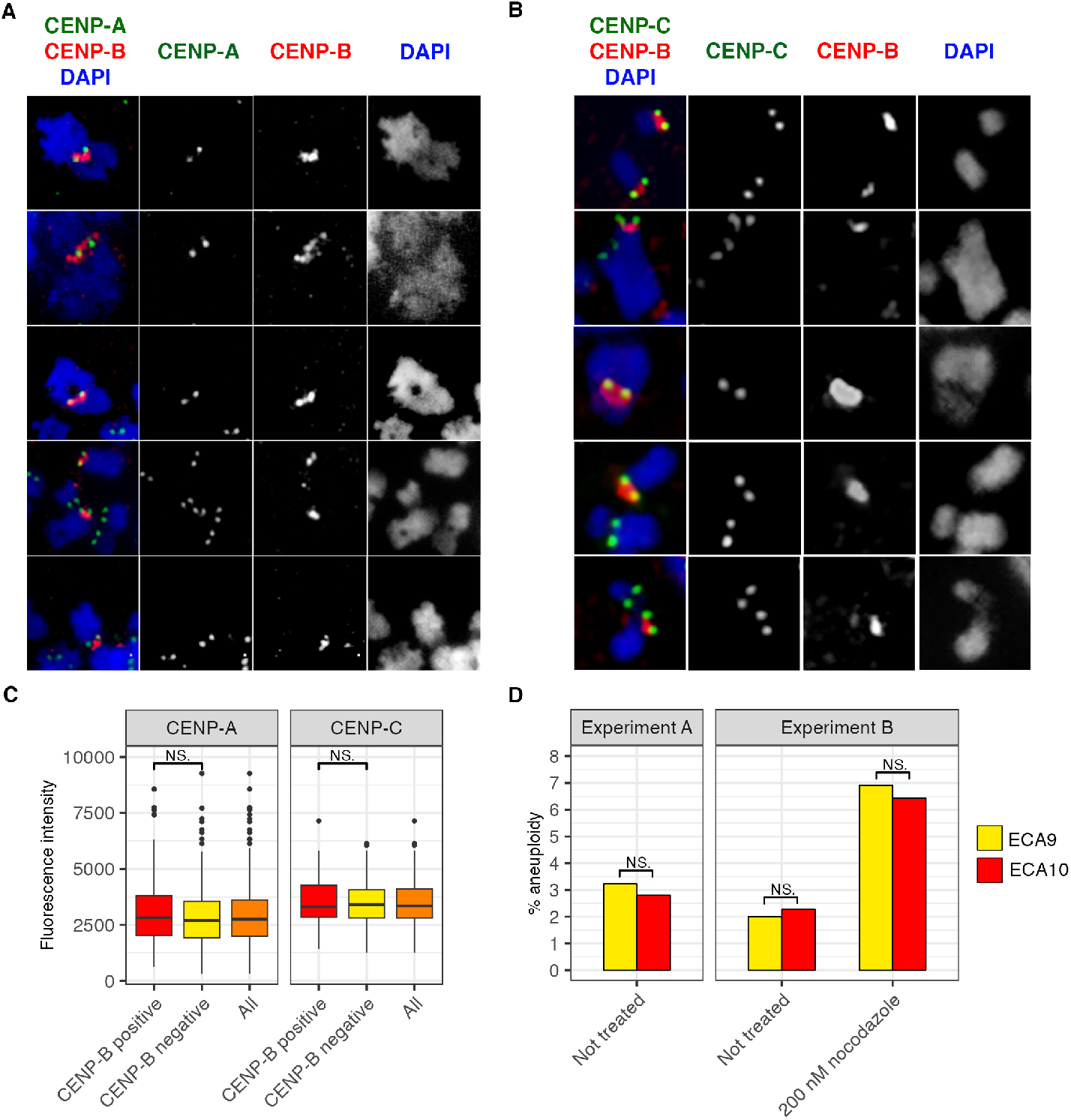
CENP-A and CENP-C localization and segregation fidelity on CENP-B positive and negative chromosomes. A) In the first column, examples of chromosomes immuno-stained with anti-CENP-B antibody (red) and anti-CENP-A serum (green). DAPI-staining is shown in blue. The three separate channels are shown in the second, third and fourth column. CENP-A signals are homogeneous both in CENP-B positive and negative centromeres. B) In the first column, examples of chromosomes immuno-stained with anti-CENP-B antibody (red) and anti-CENP-C serum (green). DAPI-staining is shown in blue. The three separate channels are shown in the second, third and fourth column. CENP-C signals are homogeneous both in CENP-B positive and negative centromeres. C) Left: CENP-A and CENP-B fluorescence intensity was measured in 96 CENP-B positive (red) and 242 CENP-B negative centromeres (yellow). Right: CENP-C and CENP-B fluorescence intensity was measured in 54 CENP-B positive (red) and 163 CENP-B negative centromeres (yellow). The values for all centromeres are reported in orange. The variability of CENP-A and CENP-C intensity is not correlated with the presence of CENP-B signals according to Student’s t test (t-value = −1.85222 and p-value = 0.064997 for CENP-A; t-value = 0.8983 and p-value = 0.185017 for CENP-C) and Spearman’s correlation test (R = 0.1125 and p-value = 0.053178 for CENP-A; R = 0.0536 and p-value = 0.432112 for CENP-C). NS: Not Significant. D) Mitotic stability of ECA9 (CENP-B negative) and ECA10 (CENP-B positive) chromosomes by interphase aneuploidy analysis. Chromosome specific BAC probes were used in FISH experiments and the number of signals per nucleus was counted in two independent experiments. Nuclei with one or three signals were considered aneuploid. In the second experiment, the number of aneuploid nuclei was counted both in normal conditions and following mitotic stress induced by a 48h treatment with 200 nM nocodazole. The numbers of counted nuclei are reported in Table S10.

We then compared the mitotic stability of a CENP-B positive (ECA10) and a CENP-B negative (ECA9) chromosome by interphase aneuploidy analysis (Fig. 5D). Chromosome specific BAC probes were used in FISH experiments and the number of signals per nucleus was counted in two independent experiments. The numbers of counted nuclei are reported in Additional file 1: Table S10. Nuclei with one or three signals were considered aneuploid. In the second experiment, the number of aneuploid nuclei was counted both in normal conditions and following mitotic stress induced by the spindle inhibitor nocodazole. The results showed that segregation fidelity was not influenced by CENP-B.

### CENP-B binding satellite and karyotype evolution

As shown in Figure 2, the CENP-B box containing satellite was detected by FISH on 21 horse, 1 donkey and 2 Grevy’s zebra primary constrictions. In addition, in Grevy’s zebra, it was detected on 16 non-centromeric ends. No signals were observed in Burchell’s zebra.

We then carried out a comparative analysis of the position of CENPB-sat loci between horse and Grevy’s zebra. In Figure 6, horse and zebra orthologous chromosomes are depicted with CENPB-sat loci indicated as red lozenges. To construct this figure, we performed a whole-genome alignment of the Equus_grevyi_HiC assembly (DNA Zoo consortium) to the horse EquCab3.0 genome (Additional file 1: Fig. S7). A detailed description of the comparative analysis is reported in the Supplementary text. Briefly, we observed four different situations: 1) maintenance of the localization of CENPB-sat at horse centromeres and at orthologous centromeric (EGR7cen/ECA15cen and EGR12cen/ECA20cen) or terminal (EGR7pter/ECA2cen, EGR8pter/ECA31cen, EGR12pter/ECA8cen and EGR16pter/ECA24cen) positions in the zebra. The terminal zebra positions can be interpreted as remnants of ancient centromeres that were inactivated in the zebra and conserved in the horse; 2) loss of CENPB-sat in the zebra compared to orthologous centromeric horse positions (EGR1/ECA25cen-ECA16cen, EGR3/ECA2q, EGR5/ECA14cen, EGR6/ECA17cen, EGR9/ECA22cen-ECA18cen, EGR11/ECA21cen-ECA19cen, EGR14/ECA6cen, EGR17/ECA10cen, EGR18/ECA8cen and EGRX/ECAX) following Robertsonian fusion or other rearrangements; 3) presence of CENPB-sat on a non-centromeric terminus of zebra chromosomes and absence on the horse orthologous chromosome (EGR2pter/ECA1pter, EGR5pter/ECA13qter, EGR13pter/ECA5pter, EGR14pter/ECA12qter, EGR15pter/ECA9pter and EGR20qter/ECA26qter). Terminal CENPB-sat zebra positions may correspond to ancestral centromeres that were inactivated in the horse; 4) presence of CENPB-sat at terminal zebra positions and at the opposite centromeric end in the horse orthologous chromosome (EGR1pter/ECA6cen, EGR6pter/ECA23cen, EGR10pter/ECA10cen, EGR19qter/ECA27cen, EGR21qter/ECA29cen and EGR22qter/ECA30cen). This peculiar comparative localization is likely a consequence of satellite DNA exchange between opposite chromosomal termini (56, 57).

**Figure 6.**
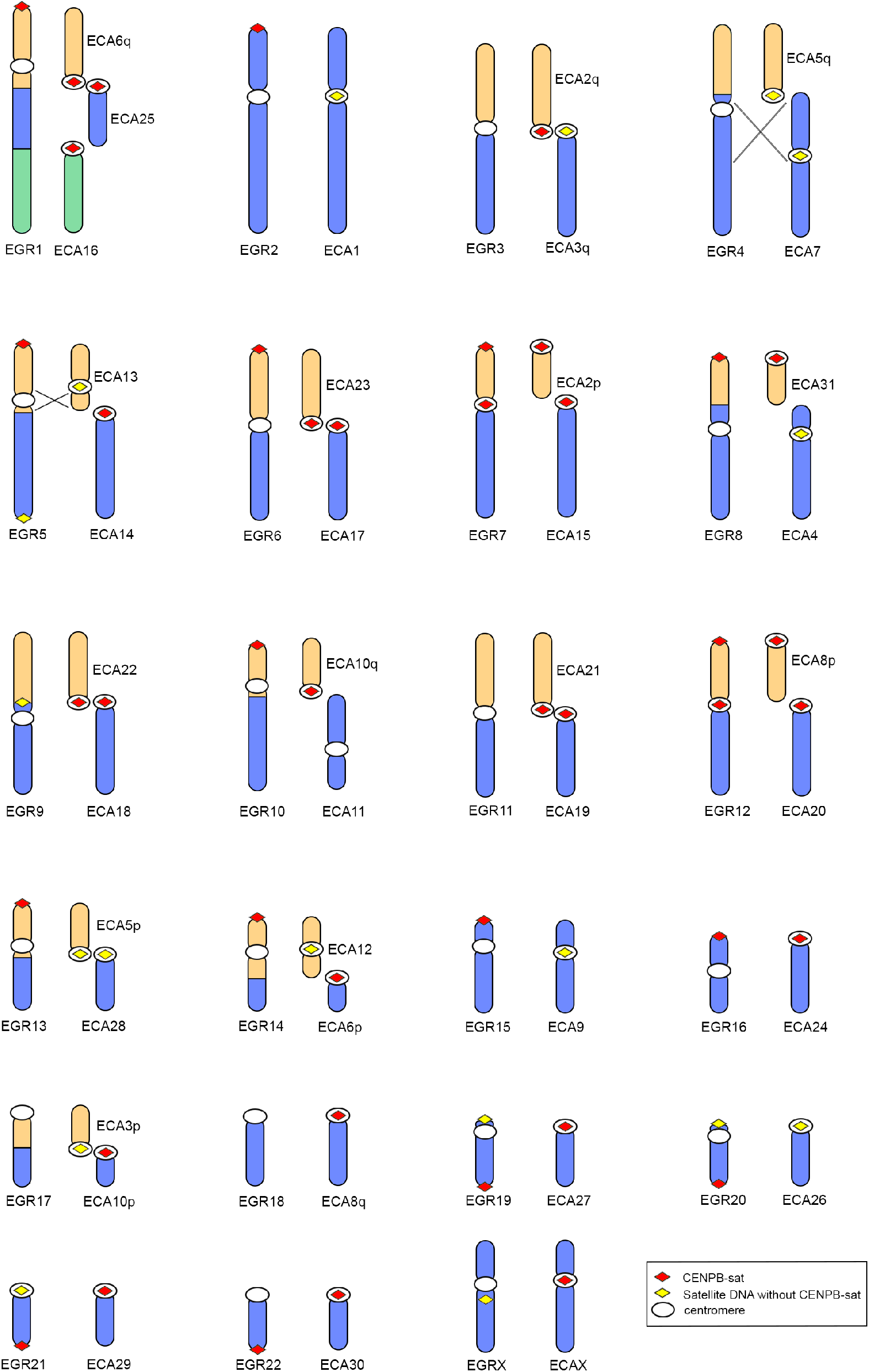
Comparison between Grevy’s zebra and orthologous horse chromosomes. Colors refer to orthologous sequences. Inverted segments are indicated with crossed lines. The position of centromeres (white ovals), CENPB-sat (red lozenges) and other satellite families (yellow lozenges) are indicated. A detailed description of this Figure is reported in Supplementary text.

## DISCUSSION

In previous work we discovered that the great karyotype heterogeneity of the otherwise closely related *Equus* species is mainly due to centromere movements that occurred during evolution through either centromere repositioning or chromosome fusion. This extensive reshuffling generated numerous satellite-free centromeres. The main question arising from these previous observations was: what makes *Equus* centromeres so plastic compared to those of the other mammalian species studied so far? In the present work, we report on a peculiarity of CENP-B binding pattern and on its dissociation from CENP-A.

The first indication that CENP-B is not associated to CENP-A came from the discovery that the 45 satellite-free centromeres that we identified in four *Equus* species do not contain any CENP-B binding motif. The CENP-B box is also missing in the main horse satellite DNA 37cen that, as we previously proved (47), is bound by CENP-A and associated to centromere competence. However, as in all mammalian species studied so far, the CENP-B protein is expressed and localized in the nucleus. Its functional domains, namely the N-terminal DNA-binding region and the C-terminal dimerization domain, are identical to those described in human and mouse, indicating that the protein is able to dimerize and bind a canonical CENP-B box.

We identified a novel equid satellite, CENPB-sat, which contains a CENP-B box and is bound by CENP-B. Although, at the cytogenetic level, CENPB-sat is localized at several primary constrictions, it is not the major centromeric satellite in any of the analyzed species but is mainly pericentromeric or located at ancestral inactivated centromeres. CENPB-sat is composed of tandemly repeated 425 bp monomers, arranged in a head-to-tail fashion. While the majority of centromeric satellites are AT rich (58), CENPB-sat is GC rich. Interestingly, we previously showed that also the major horse CENP-A binding satellite, 37cen, is GC rich (47). A 224 bp fragment of CENPB-sat, which does not contain the CENP-B box, shares 70% identity with 37cen suggesting a common evolutionary origin for CENPB-sat and 37cen.

In human and mouse, the CENP-B protein is localized at all centromeres due to the presence of CENP-B boxes within the centromeric satellites of these species. It has been stunning to observe, in the equids, a completely different binding pattern of CENP-B, which is often uncoupled from primary constrictions. In the horse, only 9 primary constrictions are bound by CENP-B while, at the unique satellite-free centromere of chromosome 11 and at 22 of the 31 satellite-based centromeres, no CENP-B binding was detected. On the other hand, the CENPB-sat satellite was detected cytogenetically at 21 primary constrictions, suggesting that, at several loci, this sequence underwent degeneration losing the ability to be recognized by the protein.

In donkey and Burchell’s zebra, CENP-B was not detected at any chromosome and the genomic amounts of CENPB-sat were extremely low. In the donkey, the consensus sequence of the CENP-B box obtained from the very few copies of CENPB-sat revealed frequent mutation of two essential nucleotides. The low enrichment of CENPB-sat in donkey CENP-B bound chromatin is another evidence of CENPB-sat degeneration. Therefore, both reduction and degeneration of binding sites are responsible for the absence of detectable levels of CENP-B. In Burchell’ zebra, the very few copies of CENPB-sat contain a canonical CENP-B box, therefore the lack of detectable CENP-B protein binding is due to the extreme paucity of binding sites.

In Grevy’s zebra the CENP-B protein was detected at two primary constrictions only and at one non-centromeric end of 13 out of the 23 chromosomes. To our knowledge, this is the first report of such extreme uncoupling between CENP-B and centromeric function. The great abundance of CENPB-sat in this species is mainly due to its localization within satellite arrays at chromosomal termini.

An intriguing finding was the presence of CENP-B enrichment peaks, identified by ChIP-seq, at intrachromosomal non-satellite positions. Several sites are shared among different species and only a subset of them contains CENP-B box-like motifs. These results suggest that CENP-B can bind DNA sequences other than the CENP-B box possibly exerting additional functions unrelated to centromeres.

In human and mouse experimental systems, it was shown that the amounts of CENP-B and CENP-C were correlated and that reduced levels of CENP-B seemed to be associated to an increased frequency of mis-segregation (14, 23, 24). On the contrary, in our natural system, the recruitment of CENP-A and CENP-C was not related to the amount of CENP-B. Indeed, while the amount of CENP-B was highly heterogeneous among different chromosomes and no detectable CENP-B was observed at several centromeres, the levels of CENP-A and CENP-C were homogeneous. These findings suggest that the interaction among CENP-B, CENP-A and CENP-C might be more complex than previously proposed.

In previous work, we compared the mitotic stability of horse chromosome 11, whose centromere is satellite-free, with horse chromosome 13, whose centromere is satellite based but, as we know now, not bound by CENP-B. We demonstrated that segregation fidelity was not influenced by the presence of satellite DNA at the centromere (59). In the present work, we compared the frequency of nuclei aneuploid for a CENP-B positive and a CENP-B negative chromosome demonstrating that mitotic segregation fidelity was not affected by the absence of CENP-B. These results are not surprising considering that CENP-B negative centromeres are fixed in the *Equus* populations and that these populations are composed by healthy, normally developing fertile individuals.

On the basis of the cytogenetic and ChIP-seq data presented in our previous (36, 47) and present work, we propose the model depicted in Figure 7 to interpret the evolution of CENPB-sat in the *Equus* species. According to this model, centromeres with CENPB-sat repeats and CENP-B binding correspond to the ancestral configuration of *Equus* centromeres which is maintained at horse chromosome 2 and Grevy’s zebra chromosome 12. In a common ancestor of all *Equus* species, the expansion of the portion of CENPB-sat lacking the CENP-B box gave rise to arrays of 37cen where the functional CENP-A binding centromere was seeded, whereas the CENP-B binding repeats were pushed towards the pericentromeric regions. Indeed, new satellite sequences are known to arise and expand in the centromeric core, progressively moving the older units towards the pericentromere, forming layers of different ages (8, 60, 61). It was proposed that pericentromeric satellites progressively become more and more degenerated and thus cannot be bound anymore by centromeric proteins, avoiding a harmful expansion of the functional centromere (9). In agreement with this view, most horse CENPB-sat loci identified by FISH are presumably degenerated and therefore no more able to bind CENP-B (Fig. 2). The presence of CENPB-sat at most horse acrocentric chromosomes (Fig. 2) further supports this hypothesis since, as mentioned above, these chromosomes correspond to ancestral ones (38, 62). Another evidence that CENPB-sat is an ancestral satellite and 37cen emerged in relatively recent evolutionary times is given by the fact that those horse metacentric chromosomes which are evolutionarily recent and derive from centromere repositioning or inversion (ECA1, ECA4, ECA7, ECA9, ECA11, ECA12, ECA13) (37) lack CENPB-sat and contain 37cen arrays. A possible explanation of this observation is that these centromeres were born satellite-free and, with the exception of the ECA11 centromere, progressively accumulated 37cen repeats during their maturation (34, 36, 41, 47). A similar situation was described in primates where centromeres deriving from centromere repositioning have acquired satellite repeats during their evolutionary maturation (48, 57).

**Figure 7.**
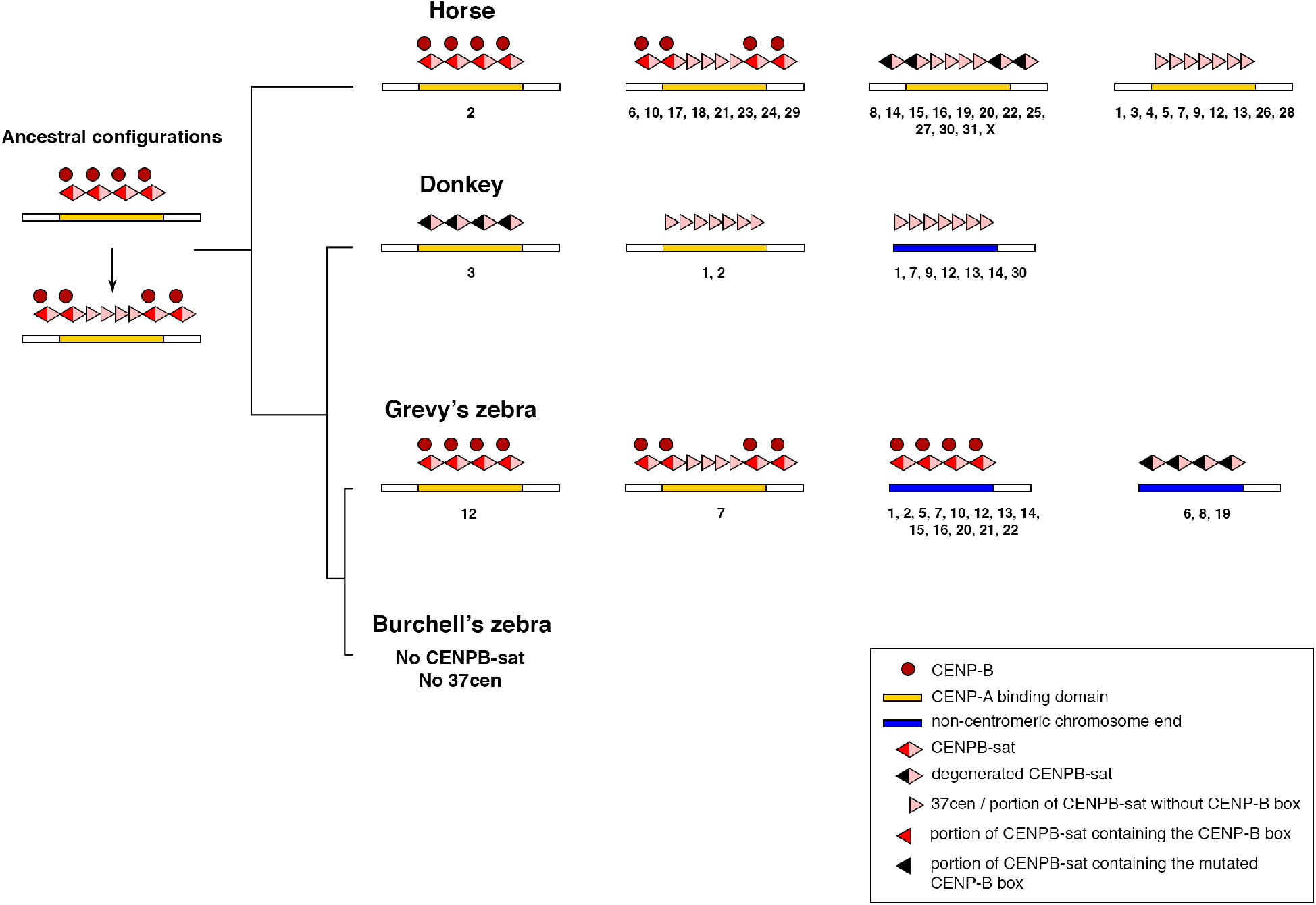
Model for CENPB-sat and 37cen evolution. The different organizations of satellites are sketched over a line representing the genomic position and CENP-A binding (yellow bars). Chromosome numbers displaying each configuration are listed under each sketch. In the equid ancestor, the centromeric CENP-A binding domains were constituted by arrays of the CENPB-sat satellite which contained a functional CENP-B box and was bound by CENP-B (red circles). Subsequently, 37cen arrays were generated by the expansion of the portion of CENPB-sat not containing the CENP-B box pushing entire CENPB-sat units outwards and colonizing the CENP-A binding domain. During the evolution of the horse lineage, at some centromeres, CENPB-sat repeats lost the ability to bind CENP-B due to mutations in the CENP-B box while at other centromeres only 37cen arrays were maintained and bound by CENP-A. In this lineage, the 37cen satellite became the major CENP-A binding centromeric satellite. In the donkey lineage, where most centromeres are satellite-free, degenerated CENPB-sat arrays were maintained at chromosome 3 only. 37cen arrays were mainly kept at non-centromeric chromosome ends (blue bars) corresponding to ancestral inactivated centromeres. In the Grevy’s zebra, the two ancestral centromere configurations can be still observed together with conserved or degenerated CENPB-sat arrays at non-centromeric chromosome ends, corresponding to inactivated centromeres. In the Burchell’s zebra, both CENPB-sat and 37cen repeats are nearly absent.

In the donkey, no binding of CENP-B was detectable and most of the few copies of the CENP-B box are degenerated. A faint CENPB-sat FISH signal at the primary constriction of chromosome 3 suggests the presence of degenerated repeats at this locus. Arrays of the 37cen satellite were observed at two donkey centromeres and at several non-centromeric chromosome ends corresponding to ancestral inactivated centromeres (Fig. 7) (36, 41).

In Grevy’s zebra, the CENPB-sat satellite is localized at numerous non-centromeric termini and at two primary constrictions (EGR12 and EGR7). In this species the non-centromeric sites correspond to inactivated centromeres or to exchange between chromosome ends (56, 57). The 37cen sequence was detected at one primary constriction only (EGR7) (36), which contains also arrays of CENPB-sat and is bound by CENP-B. The extended and conserved CENPB-sat arrays of the Grevy’s zebra at non centromeric termini could represent relics of ancestral inactivated centromeres suggesting that this species might be closer to the common ancestor than asses and other zebras. Indeed, *Equus grevyi* is the only extant member of the subgenus *Dolichohippus* and, according to paleontological, ecological and morphological evidence, is considered closer to the Eurasian ancestor than the other zebras, which are grouped in the subgenus *Hippotigris* (63, 64).

In Burchell’s zebra, we could not identify any CENP-B or CENPB-sat positive chromosome by immunofluorescence and FISH, in agreement with the extreme paucity of CENP-B binding sites revealed by sequencing. No 37cen loci were detected as well (36). In this species, the organization of satellite DNA is relatively dynamic due to the presence of novel repeats that are probably evolutionarily recent.

It is important to notice that, in all the four species, the 2PI satellite, which is not enriched in CENP-A chromatin, is one of the most abundant satellite families. This satellite is found at most horse primary constrictions, at numerous donkey and Grevy’s zebra non centromeric termini and non-centromeric interstitial or terminal positions of Burchell’s zebra (36, 44). This distribution suggests that the 2PI observed now may be the relics of the oldest equid centromeric satellite that was progressively dissociated from the centromeric function following the expansion of CENPB-sat and 37cen satellites in a lineage-specific manner. This hypothesis is supported by the variability of 2PI units (Additional file 1: Table S8 and Additional file 2: Table S9).

According to the model shown in Figure 7, in the common ancestor of equids, centromeric DNA was composed by arrays of CENPB-sat containing functional CENP-B boxes and thus binding CENP-B. A key question then arises: why, during the evolution of equids, did the centromere function escape from CENP-B binding satellite repeats landing either into arrays not containing any CENP-B box or into satellite free regions? It has been proposed that noncanonical DNA structures may contribute to centromere specification. These peculiar configurations may arise in the presence of sequence features, such as dyad symmetries or non-B DNA forming motifs, or thanks to the bending activity of sequence-specific DNA-binding proteins such as CENP-B (31). The 37cen horse sequence, as already mentioned by Kasinathan and Henikoff, is enriched in dyad symmetries which facilitate the adoption of stable secondary structures. In the present work, we found that, in the CENPB-sat sequence, dyad symmetries are restricted to the portion sharing identity with 37cen, suggesting that the expansion of 37cen satellite in the centromeric core could be favored, replacing the arrays of the entire CENPB-sat sequence in the centromeric cores. Another factor promoting the expansion of 37cen may be its length that is about half that of CENPB-sat and possibly easier to be phased with nucleosome wrapping (65–67). However, we did not find any non-B motif enrichment in the satellite-free centromeres suggesting that epigenetic factors such as heterochromatic histone marks or alterations in DNA methylation patterns may contribute to centromere specification.

In conclusion, CENP-B has been proposed to be involved in centromere strength and stability (23, 24, 68) and maintenance of pericentric heterochromatin, acting as a barrier against genome instability (27, 28). Interestingly, in donkey, Grevy’s and Burchell’s zebra, where we observed high numbers of satellite-free centromeres, karyotype reshuffling and heterogeneity in satellite DNA families, CENP-B binding was rarely observed at primary constrictions. In the horse, where only one satellite-free centromere was found, most centromeres contain CENPB-sat and a subset of them binds CENP-B. Taking together our results, we propose that the uncoupling between CENP-B and the centromeric core may drive the centromeric plasticity observed in equids. However, an important question remains open: despite being uncoupled to centromeres and poorly binding to DNA in some species, why is CENP-B well conserved and expressed in all equids? Is it simply the result of an evolutionary process or may CENP-B play extra-centromeric yet unknown roles?

## CONCLUSIONS

While in the mammalian species studied so far CENP-B and CENP-A bind the major centromeric satellite, our study showed that, in equids, CENP-B was not detectable at the numerous satellite-free and at the majority of satellite-based centromeres while it was localized at several ancestral inactivated centromeres. In horse, CENP-B binds only a fraction of primary constrictions while in donkey and Burchell’s zebra, although a functional protein is expressed, no CENP-B can be detected at centromeres. In Grevy’s zebra, CENP-B binds two centromeres only and several non-centromeric sites corresponding to ancestral inactivated centromeres. CENP-B binding sites were also detected at intra-chromosomal loci suggesting that the protein may play extra-centromeric roles.

By comparing CENP-B positive and negative centromeres, which are naturally occurring in the equid system, we demonstrated that centromeres lacking CENP-B are functional and recruit normal amounts of the centromeric proteins CENP-A and CENP-C. Thus, differently from what previously shown in human and mouse experimental systems, we proved that the role of CENP-B is more complex than previously proposed and that binding of CENP-A and CENP-C is not universally influenced by CENP-B.

The absence of CENP-B at most equid centromeres is related to the lack of CENP-B boxes rather than to peculiar features of the protein itself. While no CENP-B boxes were identified in the CENP-A binding domains of the satellite-free centromeres and in the major satellite repeat, this motif was found in a previously undescribed repeat. A comparative analysis of the localization of the CENP-B box containing satellite revealed that this satellite is an ancestral centromeric repeat which was bound by CENP-A in the common ancestor of extant equid species. We propose that, during the radiation of *Equus* species, this satellite lost the centromeric function and the resulting uncoupling between CENP-B and CENP-A may have played a role in the evolutionary reshuffling of centromeres.

These findings open a new scenario for the study of the mysterious CENP-B protein, providing new insights into the complexity of centromere organization in a largely biodiverse world where the majority of mammalian species still have to be studied.

## METHODS

### Cell lines

Primary fibroblast cell lines from horse, donkey, mule, Burchell’s zebra and Grevy’s zebra were previously described (36, 41, 44).

Primary fibroblasts were cultured in high-glucose DMEM medium, supplemented with 20% fetal bovine serum, 2 mM glutamine, 2% non-essential amino acids, 1% penicillin/streptomycin. HeLa cells were cultured in high-glucose DMEM medium, supplemented with 10% fetal bovine serum, 2 mM glutamine, 2% non-essential amino acids, 1% penicillin/streptomycin. Cells were maintained in a humidified atmosphere of 5% CO_2_ at 37°C. All the cell lines tested negative for mycoplasma.

### Antibodies

Four different commercial polyclonal anti-CENP-B antibodies were used: sc-22788 (Santa Cruz Biotechnology Inc.), raised against amino acids 535-599 mapping at the C-terminus of human CENP-B (P07199) (Fig. S1); ab84489 (Abcam), raised against a synthetic peptide corresponding to the 540-599 residues of human CENPB (P07199) (Fig. S1); the H00001059-B01P (Abnova) and the 07-735 (Sigma-Aldrich) were obtained using the entire human CENP-B protein as immunogen. Preliminary immunofluorescence experiments were carried out on human HeLa cells, horse and donkey fibroblasts to compare the three antibodies sc-22788 (Santa Cruz Biotechnology Inc.), ab84489 (Abcam), H00001059-B01P (Abnova). The results of this comparison are shown in Additional file 1: Figure S5.

Anti-CENP-A and anti-CENP-C sera were previously described (69, 70).

### ChIP-seq

Chromatin from primary fibroblasts was cross-linked with 1% formaldehyde, extracted and sonicated to obtain DNA fragments ranging from 200 to 800 bp. Immunoprecipitation was performed as previously described (41) using the anti-CENP-B sc-22788 antibody (Santa Cruz Biotechnology Inc.). Paired-end sequencing was performed with Illumina HiSeq2000 and Illumina HiSeq2500 platforms by IGA Technology Services (Udine, Italy). Reads from ChIP-seq experiments with the anti-CENP-A antibody were previously described (41, 44) and deposited in NCBI SRA Archive (SRR27325169, SRR27325168, SRR5515973, SRR5515972, SRR17956804, SRR17956803, SRR17956806, SRR17956805). The details of each dataset are reported in Additional file 1: Table S11.

### Identification of the CENP-B bound satellite, CENPB-sat, from ChIP-seq data

Reads from the ChIP-seq experiment with the anti-CENP-B antibody on horse primary fibroblasts were aligned to the horse reference genome (EquCab 2.0, 2007 release) with Bowtie (version 1.1.2), using the single end mode and *k* = 10 correction in order to refine the mapping of reads from satellite repeats (71).

Peak calling was performed using MACS14 (version 1.4.1) (72). Stringency criteria were: chrUn selection, fold enrichment > 8, -10Log_10_(p-Value) >100 and FDR (%) < 1. The 57 top-ranked regions were analyzed through Tandem Repeat Finder (73). For each region, Tandem Repeat Finder reports one or more classes of tandem repeats, providing a consensus for each class. The 425 bp consensus sequence of CENPB-sat was obtained by Multalin (74) alignment of sequences containing a canonical CENP-B box. Consensus sequences other than CENPB-sat identified by Tandem Repeat Finder were analyzed by RepeatMasker (Galaxy Version 4.1.5+galaxy0) using the RepBase library (release October 26, 2018).

To evaluate enrichment and genomic abundance of CENPB-sat in the four species, ChIP and Input reads were mapped with Bowtie2.0 (2.4.2 version) (75) using the single end mode and default parameter on the consensus sequences of the entire horse CENPB-sat or the 201 bp fragment containing the CENP-B box and not showing any identity with 37cen and ERE-1 (SAT_EC and D26566 in RepBase). Counts Per Million (CPM) from resulting BAM files were obtained using idxstats command from the Samtools package (version 1.15.1) (76). The consensus of the CENP-B box was deduced from the Input reads of each species aligned to the horse CENPB-sat sequence using the “Copy consensus sequence” function of the IGV software (2.9.2 version).

Identification of satellite repeats from unassembled Input reads was performed using TAREAN (Galaxy Version 2.3.8.1) (53) using 2 million reads as sample size. Since no satellite containing a CENP-B box was identified using Burchell’s zebra Input reads, we run the same analysis using ChIP reads obtained using the anti-CENP-B antibody. ChIP-seq mapper (Galaxy Version 0.1.1) (77) was used to evaluate the enrichment of satellite repeats identified by TAREAN in ChIP-seq experiments performed with anti-CENP-A antibody (41, 44).

### Detection of dyad symmetries and other non-B form DNA motifs

Dyad symmetries were searched in the centromeric regions we previously assembled (34, 41, 44) using EMBOSS Palindrome (version 6.6.0-7) with the minimum palindrome being 5, the maximum palindrome being 100, allowing a gap limit of 20 and allowing overlapping dyad symmetries as previously described (31, 54). Non-B DNA-forming sequence motifs, including A-phased repeats, direct repeats, inverted repeats, mirror repeats, Z-DNA and G-quadruplex, were predicted using Non-B DB v2.0 (78). For each sequence of interest, we computed the number and the coverage of sequences forming a dyad or other non-B motifs and normalized per kilobase. Centromeres containing DNA duplications (41, 44) were excluded from this analysis.

For each species, we randomly selected 100 control genomic region with a similar GC content (37±1.5 for the horse, 35±1.5 for the donkey, 36.6±1.5 for the Grevy’s zebra and 37±1.5 for the Burchell’s zebra; these ranges correspond to the average GC ± the standard deviation of the GC content of the centromeric regions) and a length corresponding to the average length of the centromeric domains (500 kb for the horse, 404 kb for the donkey, 237 kb for the Grevy’s zebra and 225 kb for the Burchell’s zebra). The selection of control regions was performed using bedtools (v2.30.0) and Seqkit (v2.6.1). Reference genomes used to identify control regions were the horse EquCab2.0 assembly (34), the donkey ASM1607732v2 assembly, the Grevy’s zebra Equus_grevyi_HiC assembly (49) and the Burchell’s zebra Equus_quagga_HiC assembly (49).

We calculated standardized Z-score for the values of the unique horse satellite-free centromere (ECA11) with respect to control regions. To test whether the differences were statistically significant, we calculated the P-values using Z-score calculator (79). For the other species, in case of normal distribution, unpaired, two-tailed t-test or unpaired two-tailed Welch’s t-test were used (80). In the case of non-Gaussian distribution, two-tailed Mann-Whitney U test was applied (79). Boxplots with statistical significance analysis were obtained using ggplot2 and ggsignif R packages.

### Genome-wide analysis of CENP-B binding sites

To evaluate the genome-wide distribution of CENP-B binding sites, ChIP-seq reads from the four species were aligned with paired-end mode to the EquCab2.0 reference genome with Bowtie2 (version 2.4.2) using default parameters (71, 75). Peak calling was performed with MACS2 (version 2.2.7.1) (72) using 0.01 as q-value cutoff. We excluded from the analysis the peaks overlapping satellite sequences using UCSC Table Browser and the peaks identified in unplaced contigs. CENP-B boxes were searched using FIMO (81). The content of interspersed repeats was analyzed with RepeatMasker using the RepBase library (release October 26, 2018). Bedtools (v2.30.0) was utilized to identify peaks shared among different species.

### Sequencing of *CENP-B* genes

The sequence of the *CENP-B* coding sequence of horse, donkey, Grevy’s zebra and Burchell’s zebra was obtained by Sanger sequencing of PCR fragments and by directly assembling reads from ChIP-seq input datasets. Primers used for PCR amplification and sequencing are listed in Additional file 1: Table S12.

### Western blotting

Total protein extracts were prepared from samples of three million cells as follows: the cells were washed twice with ice cold 1xPBS, resuspended in lysis buffer (50 mM Tris–HCl pH 6.8, 86 mM β-mercaptoethanol, 2 % SDS) and boiled for 10 min, as previously described (52). Nuclear and cytoplasmic protein extracts were prepared using the fractionation protocol developed by Suzuki and colleagues (82). Briefly, starting from samples of 30 million cells, the cells were resuspended in ice-cold 0.1% NP40 in PBS. The nuclear and cytoplasmic fractions were then separated by a 10 second centrifugation at 6000 rpm. The supernatant was saved as cytoplasmic fraction, diluted in Laemmli buffer and boiled for 1 minute. The pellet was resuspended in ice-cold 0.1% NP40 in PBS, centrifuged again as above and the final pellet was resuspended in Laemmli buffer, sonicated, boiled for 1 minute and saved as nuclear extract.

Proteins were separated by SDS-PAGE on polyacrilamide gel and blotted to nitrocellulose membranes (Amersham^TM^ Hybond^TM^-ECL, GE-Healthcare) according to standard methods. Membranes were incubated with the anti-α tubulin antibody [DM1A] ab7291 (Abcam), diluted 1:5000, the anti-CENP-B sc-22788 antibody (Santa Cruz Biotechnology Inc.), diluted 1:750 or with the anti-CENP-B 07-735 antibody (Sigma-Aldrich), diluted 1:1000. HRP conjugated secondary antibodies were used. Pre-incubation of membranes and dilutions of antibodies were performed in 1x PBS containing 0.05% Tween-20 and 7.5% skim milk. Detection was performed using the BioRad Clarity^TM^ Western ECL Substrate kit following manufacturer’s procedures.

### CENPB-sat plasmid vector construction

The portion of the CENPB-sat comprising the CENP-B box and lacking identity regions with the 37cen satellite was amplified from horse genomic DNA using the following primer oligonucleotides containing EcoRI and SalI adapters required for cloning purposes: CENPBsat-F 5’-ATTGAATTCCCTTTCTGACATAGGTGCTTTCTG-3’ and CENPBsat-R 5’-ATTGTCGACGCTTTAGGACTTCTGCTTCTG-3’. PCR products were digested with EcoRI/SalI and cloned in the pSVal plasmid (83). An 8-copies array of the cloned portion was obtained as previously described (50).

### Immunofluorescence and FISH

We carried out preliminary immunofluorescence experiments to test several permeabilization and fixation procedures with the three anti-CENP-B antibodies described above (sc-22788 Santa Cruz Biotechnology Inc., ab84489 Abcam or H00001059-B01P Abnova). The best combination was fixation with ice-cold methanol for 4 minutes followed by permeabilization with 1x PBS 0.05% Tween-20 for 15 minutes at room temperature and incubation at 37°C for 2 hours with H00001059-B01P Abnova antibody diluted 1:100. Incubation with the anti-CENP-A (70) or anti-CENP-C serum (69), both diluted 1:100, was carried out at 37°C for 1 hour. Digital grey-scale images were acquired with a fluorescence microscope (Zeiss Axio Scope.A1) equipped with a cooled CCD camera (Photometrics) using a 63x oil objective. In immuno-FISH experiments, immunofluorescence signals were collected before hybridization with the CENPB-sat FISH probe. Pseudo-coloring and merging of images were performed using the IpLab software.

Metaphase spreads were obtained with the standard air-drying procedure. CENPB-sat plasmid extraction, nick translation with Cy3-dUTP (ENZ-42501) and hybridization were performed as previously described (36). Whole genomic DNA was used as probe for total satellite DNA. Chromosomes were counterstained with DAPI and identified by computer-generated reverse DAPI banding according to the published karyotypes.

3D-immunofluorescence on whole cells was performed using a slight modification of the protocol described by Solovei and Cremer (84). Cells were grown on coverslips, rinsed with PBS and fixed with 4% paraformaldehyde in PBS at room temperature. During the last minute of fixation, a few drops of 1x PBS 0.5% Triton X-100/PBS were added. After three washes in 0.01% Tween-20, cells were permeabilized with 1x PBS 0.5% Tween-20 for 20 minutes at room temperature. Anti-CENP-B (sc-22788 Santa Cruz Biotechnology Inc.) and anti-tubulin (ab7291 Abcam) antibodies were diluted 1:80 and 1:500, respectively. Stacks of optical sections through whole cells were collected using a Leica TCS SP8 STED 3X confocal microscope (Centro Grandi Strumenti, University of Pavia).

### Quantification of immunofluorescence signals

Quantification of CENP-A, CENP-B and CENP-C signal intensities on metaphase spreads was performed using Fiji software (85). The integrated signal density of each centromeric signal was calculated by subtracting the fluorescence intensity of the background from the total intensity of the signal. Statistical significance was evaluated using Spearman’s Rho correlation test and unpaired two-tailed t-test (79). Boxplots with statistical significance analysis were obtained using ggplot2 and ggsignif R packages.

### CENP-B-eGFP plasmid construction and transfection

The horse *CENP-B* coding sequence was cloned upstream of the enhanced Green Fluorescent Protein (eGFP) cDNA, into an expression vector that was previously constructed in our laboratory (83). The vector contains the puromycin resistance genes. The chimeric protein was expressed under the control of the Cytomegalovirus-immediate early (CMVie) promoter.

The plasmid (pCCB-GFP) was used to transfect a horse fibroblast cell line, previously immortalized in our laboratory (55). Transfection was carried out using the Neon™ Transfection System (Thermo Fisher Scientific) according to the manufacturer’s protocol. 48 hours after transfection, puromycin (750 ng/ml) was added and resistant clones were isolated after three weeks. Cells were then harvested by trypsinization, treated with hypothonic 75mM KCl solution for 25 minutes at 37°C, cyto-spun onto slides at 1250 rpm for 8 minutes and fixed with ice-cold methanol for 4 minutes.

### Interphase aneuploidy analysis

For each experiment, 3×10^5^ cells were seeded in 10 cm plates. Untreated cells were grown for 72 hours, while treated cells were exposed to 200 nM nocodazole (Sigma-Aldrich) after 24 hours culture period and then grown for the remaining 48 hours. Cells were then harvested by trypsinization, treated with hypothonic solution (75mM KCl) for 25 minutes at 37°C and then fixed with cold 1:3 acetic acid:methanol solution over-night at 4°C. Nuclei were then spread onto slides according to the standard air-drying procedure.

To identify horse chromosomes 9 and 10, two bacterial artificial chromosomes derived from the CHORI-241 BAC library (CH241-361E21, chr9:36,816,509-36,983,616 in EquCab3.0; CH241-403K5, chr10:28,469,205-28,662,312 in EquCab3.0) were extracted from 10 ml bacterial cultures with the Quantum Prep Plasmid miniprep kit (BioRad), according to supplier instructions. The probes were labeled by nick translation with Cy3-dUTP (Enzo Life Sciences) and FISH was performed as previously described (36). The χ^2^ test was used to evaluate whether the differences in the frequency of aneuploid nuclei were statistically significant.

## DECLARATIONS

### Ethics approval and consent to participate

Not applicable

### Consent for publication

Not applicable

### Availability of data and materials

Raw sequencing data from this study are available in the NCBI BioProject database (https://www.ncbi.nlm.nih.gov/bioproject/) under accession number PRJNA1054998. In this work we also used publicly available ChIP-seq datasets (SRR27325169, SRR27325168, SRR5515973, SRR5515972, SRR17956804, SRR17956803, SRR17956806, SRR17956805) that we previously deposited in NCBI SRA Archive.

### Authors’ contributions

EC and FMP carried out most molecular and cell biology experiments and bioinformatic analyses. SGN, MB, EG and ER contributed to some molecular and cell biology experiments. EG, EC and FMP conceived the study and wrote the manuscript. EG supervised the study. All authors participated in discussions and result interpretation. All authors read and approved the final manuscript.

## Acknowledgements

We would like to thank Terje Raudsepp (Texas A&M University, USA) for providing to us the horse BAC clones, Douglas F. Antczak and Donald Miller (Cornell University, USA) for the mule fibroblast cell line, Sergio Comincini (University of Pavia) for helpful suggestions on cell transfection by electroporation, Giulio Pavesi (University of Milan), Riccardo Gamba and Kevin Sullivan (University of Galway, Ireland) for advice on bioinformatic analyses and plasmid construction.

## LIST OF SUPPLEMENTARY FILES

**Additional file 1**: PDF, Supplementary Materials.

This file includes Supplementary Text, Supplementary Figures S1 to S7, Supplementary Tables S1, S3, S4, S5, S8, S10, S11 and S12.

**Additional file 2**: XLSX, Supplementary Tables_2_6_7_9. This file includes Supplementary Tables S2, S6, S7 and S9.

**Additional file 3**: AVI, Movie S1 (related to Figure 4)

3D fluorescence imaging of CENP-B (red) and tubulin (green) in HeLa cells. Arrows indicate the cells shown in Figure 4.

**Additional file 4**: AVI, Movie S2 (related to Figure 4)

3D fluorescence imaging of CENP-B (red) and tubulin (green) in horse primary fibroblasts. Arrows indicate the cells shown in Figure 4.

**Additional file 5**: AVI, Movie S3 (related to Figure 4)

3D fluorescence imaging of CENP-B (red) and tubulin (green) in horse primary fibroblasts. The arrow indicates the cell shown in Figure 4.

**Additional file 6**: AVI, Movie S4 (related to Figure 4)

3D fluorescence imaging of CENP-B (red) and tubulin (green) in Grevy’s zebra primary fibroblasts. Arrows indicate the cells shown in Figure 4.

**Additional file 7**: AVI, Movie S5 (related to Figure 4)

3D fluorescence imaging of CENP-B (red) and tubulin (green) in donkey primary fibroblasts. Arrows indicate the cells shown in Figure 4.

**Additional file 8**: AVI, Movie S6 (related to Figure 4)

3D fluorescence imaging of CENP-B (red) and tubulin (green) in donkey primary fibroblasts. Arrows indicate the cells shown in Figure 4.

**Additional file 9**: AVI, Movie S7 (related to Figure 4)

3D fluorescence imaging of CENP-B (red) and tubulin (green) in Burchell’s zebra primary fibroblasts. Arrows indicate the cells shown in Figure 4.

**Additional file 10**: AVI, Movie S8 (related to Figure 4)

3D fluorescence imaging of CENP-B (red) and tubulin (green) in Burchell’s zebra primary fibroblasts. Arrows indicate the cells shown in Figure 4.

