## Additional file 1 for "CENP-A/CENP-B uncoupling in the evolutionary reshuffling of centromeres"

**This PDF file includes:**

Supplementary Text
Figs. S1 to S7
Tables S1, S3, S4, S5, S8, S10, S11; S12

**Other Supplementary Materials for this manuscript include the following:**

Additional file 2: Supplementary excel file containing Tables S2, S6, S7, S9
Additional files 3-10: Movies S1 to S8

**Supplementary Text**

Comparative analysis of CENPB-sat cytogenetic localization in Grevy's zebra and horse chromosomes

EGR1 is a metacentric chromosome deriving from the fusion of three ancestral acrocentric elements that correspond to ECA6q, ECA25 and ECA16. In the horse, ECA6, ECA15 and ECA25 display CENPB-sat at their primary constriction. However, all the ancestral satellite loci were lost during the fusion events that originated the EGR chromosomes, which carries a satellite-free centromere far from the ECA6q/ECA25 fusion site (1). The CENPB-sat satellite is present at the p-terminus of the EGR chromosome.

EGR2 is a metacentric chromosome orthologous to ECA1. While in EGR2 the CENPB-sat is present at the p-terminus, in the horse no CENPB-sat loci were detected and other satellite families were previously observed at the primary constriction of this chromosome (2). No satellite sequences were detected at the primary constriction of EGR2 at the cytogenetic level (2). However, since no satellite-free centromere was identified (1), we cannot exclude that short or chromosome-specific tandem arrays, undetectable at the FISH resolution level, might be present. Interestingly, ECA1 probably does not represent the ancestral configuration, since a fusion event occurred in the equid ancestor (3, 4). Thus, the CENPB-sat locus at EGR2pter may correspond to an ancestral centromere that was inactivated during equid evolution.

EGR3 chromosome is orthologous to ECA2q and ECA3q. Interestingly, the syntenic group ECA 2q+3q is maintained in all equid species with the exception of the horse, suggesting that, in this case, the horse does not carry the ancestral configuration (3-6). On the EGR3 chromosome, no satellite loci were detected by FISH and its centromere is satellite-free (1, 2). However, it is worth noticing that the donkey ortholog (EAS3) is the unique donkey chromosome carrying a CENPB-sat locus, suggesting that it may represent the ancestral configuration. In the horse, chromosome ECA2 only shows CENPB-sat at its primary constriction.

EGR4 derived from the fusion of elements orthologous to ECA5q and ECA7. In both species, no CENPB-sat loci were identified. In the zebra chromosome no satellite arrays were detected and the centromere is satellite-free (1), while, in the horse, ECA5 and ECA7 chromosomes carry other satellite families at their primary constrictions (2). Crossed lines indicate inverted segments (Figure S7).

EGR5 was originated by the fusion between ancestral elements corresponding to ECA13 and ECA14. While ECA14 carries CENPB-sat at its centromeric terminus, satellite sequences were lost during the fusion generating a satellite-free centromere (1). Interestingly, EGR5 carries CENPB-sat at its p-terminus which corresponds to the q-terminus of ECA13. According to previous cytogenetic results, the metacentric ECA13 does not represent the ancestral configuration which is supposed to be acrocentric (3). These observations suggest that the CENPB-sat locus at EGR5pter may correspond to the inactivated centromere of the ancestral acrocentric element. Crossed lines indicate inverted segments (Figure S7).

EGR6 derives from the centric fusion of the ancestral acrocentrics orthologous to ECA23 and ECA17, which both carry CENPB-sat at their primary constrictions. At the fusion region, satellite sequences were lost during the formation of its satellite-free centromere (2), while the CENPB-sat locus observed at EGR6pter is likely due to exchange between chromosome ends.

EGR7 was originated from fusion between the ancestral acrocentrics orthologous to ECA2p and ECA15. During this fusion, all CENPB-sat loci were maintained.

EGR8 derives from the fusion of elements orthologous to ECA31, which carries CENPB-sat at its centromere, and ECA4, which displays satellite sequences other than CENPB-sat. EGR8 shows CENPB-sat at its p-terminus in correspondence to the ECA31 centromeric locus. No satellite sequences were detected at the primary constriction of EGR8 at the cytogenetic level (2). However, since no satellite-free centromere was identified by ChIP-seq (1), it is likely that short or chromosome-specific tandem arrays, undetectable at the FISH resolution level, are present.
EGR9 is orthologous to ECA22 and ECA18, which both carry CENPB-sat and other satellite repeats (2) at their primary constrictions. However, the zebra chromosome displays only satellite sequences other than CENPB-sat at the fusion region, suggesting that CENPB-sat arrays were lost during the fusion. On EGR9, the presence of a satellite-free centromere was demonstrated by ChIP-seq (1). As previously discussed (1), the centromeric function moved away from satellite repeats.
EGR10 derives from the fusion of ancestral elements orthologous to ECA10q and ECA11. At the fusion site, satellite arrays were lost but CENPB-sat is present at the p-terminus of EGR10, which correspond to the q-terminus of ECA10. The CENPB-sat locus observed at EGR10pter is likely due to exchange between chromosome ends.
EGR11 was originated from a centric fusion between two elements corresponding to ECA21 and ECA19. In the horse, both chromosomes carry CENPB-sat at their primary constrictions, while in the Grevy's zebra all satellite arrays were lost during the fusion (1).
The situation is different in EGR12, that originated by the fusion of two ancestral acrocentrics corresponding to ECA8p and ECA20. As for EGR7, Grevy's zebra and horse show CENPB-sat loci at orthologous positions suggesting that ancestral satellite arrays were maintained.
EGR13 originated from the fusion of ancestral elements that correspond to ECA5p and ECA28. In the zebra chromosome, no satellite sequences were detected at the fusion region while a CENPB-sat locus is present at the p-terminus of the chromosome.
EGR14, orthologous to ECA12 and ECA6p, shows a CENPB-sat locus at its p-terminus. The metacentric configuration of ECA12 does not correspond to the ancestral configuration which is supposed to be acrocentric (3), suggesting that the CENPB-sat locus at EGR14pter may correspond to the inactivated centromere of the ancestral acrocentric.
EGR15 is entirely colinear to ECA9. Similar to what described for ECA13 and ECA12, we hypothesize that the ancestral element was acrocentric and that the CENPB-sat locus at EGR15pter may correspond to the ancestral inactivated centromere.
The centromere of EGR16 is satellite-free and derives from a centromere repositioning event (1). The orthologous horse ECA24 retains the ancestral acrocentric configuration with CENPB-sat at its primary constriction. CENPB-sat is still detectable at EGR16pter, as remnant of the ancient inactivated centromere.
EGR17 derived from fusion between the ancestral elements corresponding to ECA3p and ECA10p. No satellite sequences were detected at the primary constriction of EGR17 at the cytogenetic level (2). However, since no satellite-free centromere was identified by ChIP-seq (1), it is likely that short or chromosome-specific tandem arrays, undetectable at the FISH resolution level, are present.
EGR18 is orthologous to ECA8p. While CENPB-sat repeats were detected at the primary constriction of ECA8 no satellite repeats were identified by FISH on EGR18.
In our previous work, we showed that the centromeres of EGR19 and EGR20 derived from repositioning and are satellite-free. The orthologous of EGR19 is the acrocentric ECA27 which carries CENPB-sat at the primary constriction while the orthologous of EGR20 is the acrocentric

ECA26 which carries other satellites at the primary constriction (2). An exchange between termini may have occurred in the zebra moving CENPB-sat repeats at the q terminus. A similar situation was observed in EGR21 and EGR22 which are colinear with ECA29 and ECA20, respectively. However, at these chromosomes, ChIP-seq did not detect satellite-free centromeres (1).
EGRX is entirely colinear with ECAX, which shows CENPB-sat at its primary constriction. However, ChIP-seq experiments showed that the zebra chromosome carries a satellite-free centromere (1).

**Fig. S1**

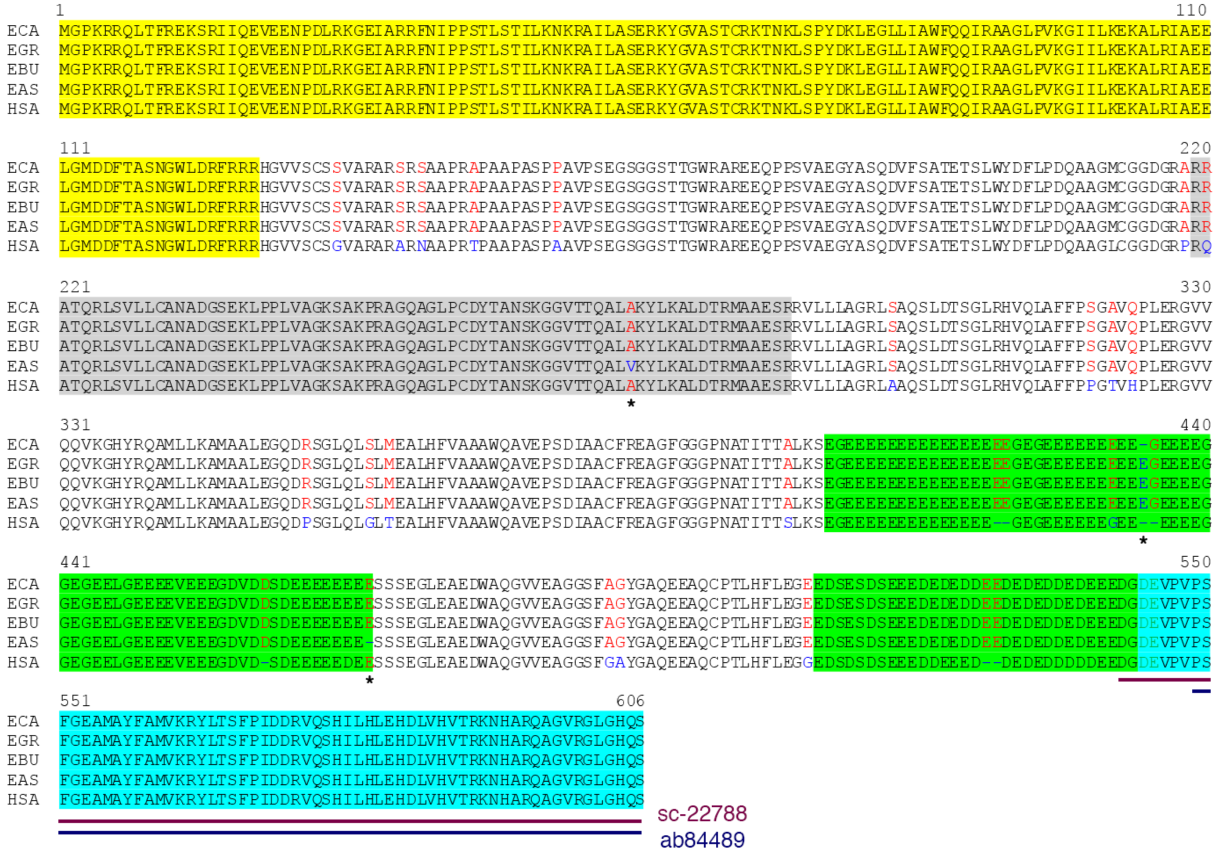


**Figure S1. Alignment of horse (ECA), Grevy's zebra (EGR), Burchell's zebra (EBU), donkey (EAS) and human (HSA) CENP-B proteins.** The human protein NP_001801.1 was used as reference. Protein sequences from the other species were deduced from genomic sequences. The DNA binding domain, the endonuclease domain, the acidic domains and the dimerization domain are highlighted in yellow, grey, green and turquoise blue, respectively. Amino acid differences are shown using blue (high consensus color) or red (low consensus color). The three amino acid differences among the equid CENP-B proteins are marked with an asterisk. The epitopes of anti-CENP-B sc-22788 (Santa Cruz Biotechnology Inc.) and anti-CENP-B ab84489 (Abcam) are shown.

### Fig. S2

**A**

```
                   1                                                                                                  100
CENPB-sat-201-425  GTTAAGCGCTGAAAAGAAATGGCNTTTCAGCTGCCTTTGTATGAGATGTTCCCAGGNACGCTGTAAGAGCACTGTGGAAAGCGAGTTCTTTCTCAGCTTC
            37cen  GATCAGGCCTGCAAAGAAACTGCGTTTCACAGGCCTTTGGAAGAGATGTTCCC-GGTAGGCTGTAAGAGCACTGTGCAGAGCGAGTTGTTTCTTAGCTTC

                   101                                                                                               199
CENPB-sat-201-425  CTAAAGAGCTGGANGGCAAGACAGTTTATGGCTTGTCTCCCATTGAAGGA-TGGAGGCAGTGCTTTGTGCCTTCCACCTCTAGAGCAATGGAGGGCACGG
            37cen  CCAAAGAGCTGGAAG-CAAGATGCTGTGGGGCCCAACTCGCCCTTT-GGAAAGAAGCCTGCACGTTGTGCCTTTCAGCTCTAGGGCAAAGTAGCACACCC

                   200                      225
CENPB-sat-201-425  CTGAG-AGCAAAGGGGCCTTTCTGACA
            37cen  A-GAGCAGAAGTCCTAC-TT-CAGCCA
```

**B**

```
               1                                                                                                  100
  CENPB-sat    TAGGTGCTTTCTGACACTCTCTNNANCCAGTGCACAATGGTGGTTTGTAAAAAGCCTATTGTCTGNCTCCTCCTAAGCATGTGGAAGCACAATCATTTGG
 EGR-Tarean    TAGGTGCATTCTGACACTCTCTGA-CCCAGTGCACCATGGCGGTTTGTAAAAAGCCTATTGTCTGC-TCCTGATAAGCATGTGGAAGCACAGTCTTCTGG
 EBU-Tarean                    CTCTCTCCACC-AGTGCACAATGGTGGTTGGAAAAAGGCGTGCAGTCTATCGCCTCCTAAGCATGTGGAAGCTCAATCATTTGG
 EAS-Tarean    TAGGCGTTTTCTGACACTCTCTCCACC-AGTGGTCAACTGTGGTTGGTGAAAAGCGTATTGTCTCTCTCCTGCTAAACACATGGTGGCACAATCATGTGG

               101                                                                                                200
  CENPB-sat    GCCTCGNCCCCGTTGCTTAAGGGGAAGATGTAGGCATTTCGTCTGAGCCGGGTTGGCAAGGTGGAAATGTGGCAGAAGCAGAAGTNCCAAAGCTNGNCGA
 EGR-Tarean    GCCTCGCCCCCATTGCTTAAGGGGAAGATGTAGGCATTTCGTCTGAGCCGGGTTGGCAAGGGGAAATCTGGCAGAAGCAGAAGTTCTAAAGC--GGCGA
 EBU-Tarean    GACTCAGCCCAGTTGCTTAAGGGGAAGATGTAGGCATTTCGTCTGAGCCGGGTTGGCAAGGTGGAATTGTGGCAGAAGCAGAAGTTCTAAAGC--GGCGA
 EAS-Tarean    GACTCAGCCCAGTTGCTTAAGGGGAAGATGTAGGCATTTTGTCTGAGCTGGGTTGGCAAGGTGGAACGGCGGCAGAAGCGGAAGTTCTAAAGC--GGCGA

               201                                                                                                300
  CENPB-sat    GTTAAGCGCTGAAAAGAAATGGCNTTTCAGCTGCCTTTGTATGAGATGTTCCCAGGNACGCTGTAAGAGCACTGTGGAAAGCGAGTTCTTTCTCAGCTTC
 EGR-Tarean    GTTAAGCGCTGGGAAGAAATGGCATTTGAGCTGCCTTTGTATGAGATGTTCCCAG-TATGCTGTAACAGCACTGTTGAAAGCGAGTTCTTTCTCAGCTTC
 EBU-Tarean    GTTAAGGGCTGGGAAGAAATGGCGTTTGAGCTGCCTTTGTATGAGATGTTCCCAG-TATGCTGTAAGAGCACTGTGGAAAGCGAGTTCTTTCTCAGCTTC
 EAS-Tarean    GTTAAGGGCTGGGAAGAAATGCCACTGGAGCTGCCTTTGTATGAGAAGTTCCC-GGTCTGCTGTAACAGGACTGTGGCAAGCGAGTTGTTTGTCAGCTTC

               301                                                                                                400
  CENPB-sat    CTAAAGAGCTGGANGGCAAGACAGTTTATGGCTTGTCTCCCATTGAAGGATGGAGGCAGTGCTTTGTGCCTTCCACCTCTAGAGCAATGGAGGGCACGGC
 EGR-Tarean    CTAAAGAGCTGGAGG-CAAGACAGTTTATGGCTTGTCTCCCATTGAAGGATGGAGGCAGTGCTTTGTGCCTTCCACCTCTAGAGCAATGGAGGGCACGGC
 EBU-Tarean    CTAAAGAGCTGGAGG-CAAGACAGTTTATGGCTTGTCTCCCATTGAAGGATGGAGGCAGTGCTTTGTGCCTTCCACCTCTAGAGCAATGGAGGGCACGGC
 EAS-Tarean    GTAAAGGGCTGGAGG-CAAGACAGTTTATGGCGTGTCTCCCATTGAAGGATGGAGGCAGTGCTTTGTGCCTTCCACGTCTAGAGCAATGGAGGGCACGGC

               401                      425
  CENPB-sat    TGAGAGCAAAGGGGCCTTTCTGACA
 EGR-Tarean    TGAGAGCAAAGGGCCCTTTCTGACA
 EBU-Tarean    TGAGAGCAAAGGGGCCTTTCTGACA
 EAS-Tarean    TGAGAGCAAAGGGGCCTTTCTGACA
```

**Figure S2. CENPB-sat sequence. (A)** Alignment between the 201-425 nt region of the CENPB-sat and 37cen satellite. High and low consensus nucleotides are colored in red and blue, respectively. **(B)** Alignment of horse CENPB-sat consensus with the consensus sequences of CENPB-sat of Grevy's zebra (EGR), Burchell's zebra (EBU) and donkey (EAS) obtained from input reads using TAREAN. High and low consensus nucleotides are colored in red and blue, respectively. CENP-B boxes are highlighted in yellow. The nucleotides essential for CENP-B binding are underlined.

### Fig. S3

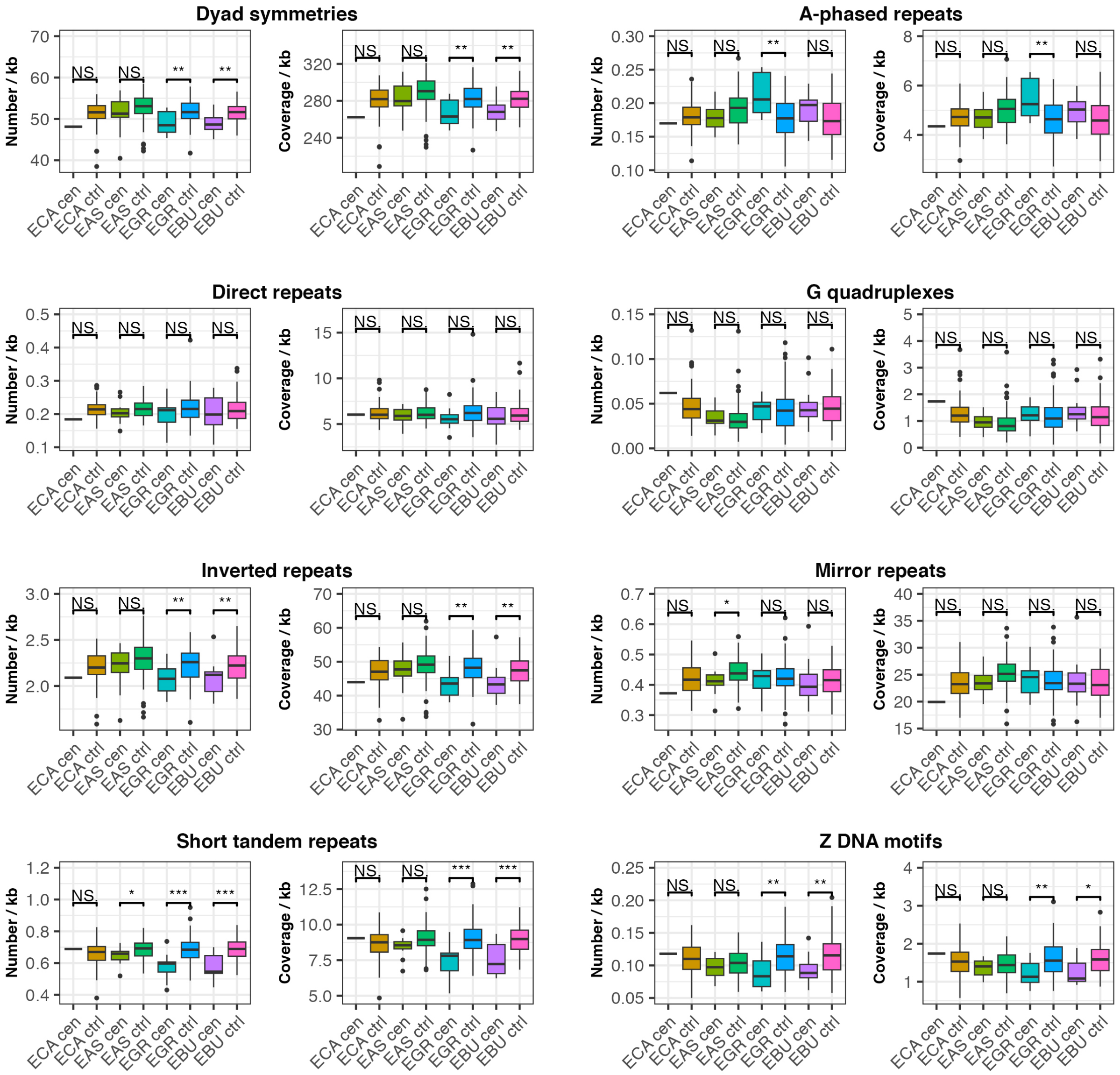


**Figure S3. Non-B DNA forming motifs in satellite-free centromeric and control regions.** For each type of motif, boxplots of normalized numbers and coverage are shown. Genomic regions with the same length and AT content of centromeric regions were used as control (ctrl). Dyad symmetries were searched using EMBOSS palindrome. A-phased repeats (bent DNA), direct repeats (slipped DNA), G quadruplexes, inverted repeats (cruciform DNA), mirror repeats (triple-helix DNA), short tandem repeats and Z-DNA were searched using Non-B DB v2.0. Levels of significance are reported with asterisks.

**Fig. S4**

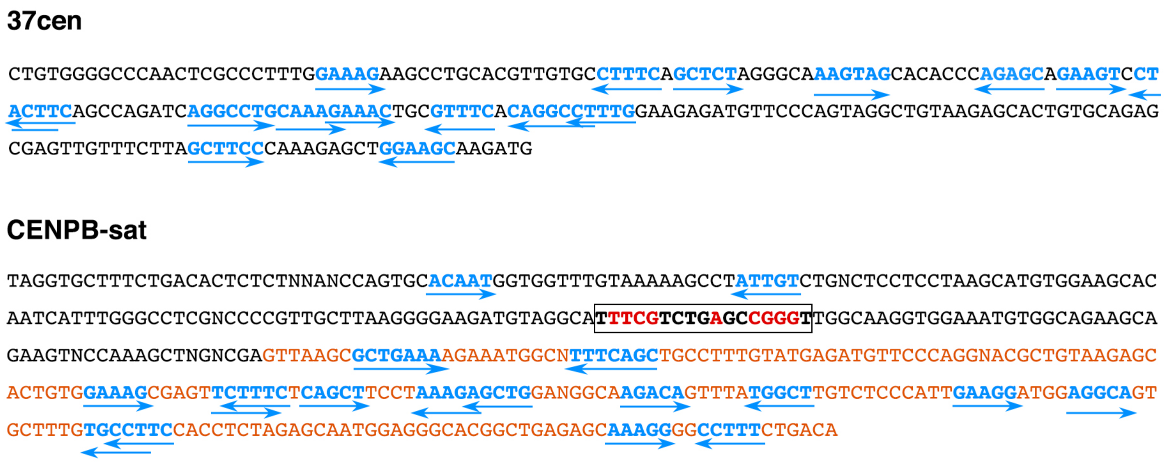


**Figure S4. Dyad symmetries in the consensus sequences of horse CENPB-sat and 37cen satellites.** Sequences forming dyads are colored in cyan and indicated with arrows. In the CENPB-sat sequence, a rectangle indicates the CENP-B box with the essential nucleotides written in red. The portion sharing identity with 37cen sequence is written in orange.

**Fig. S5**

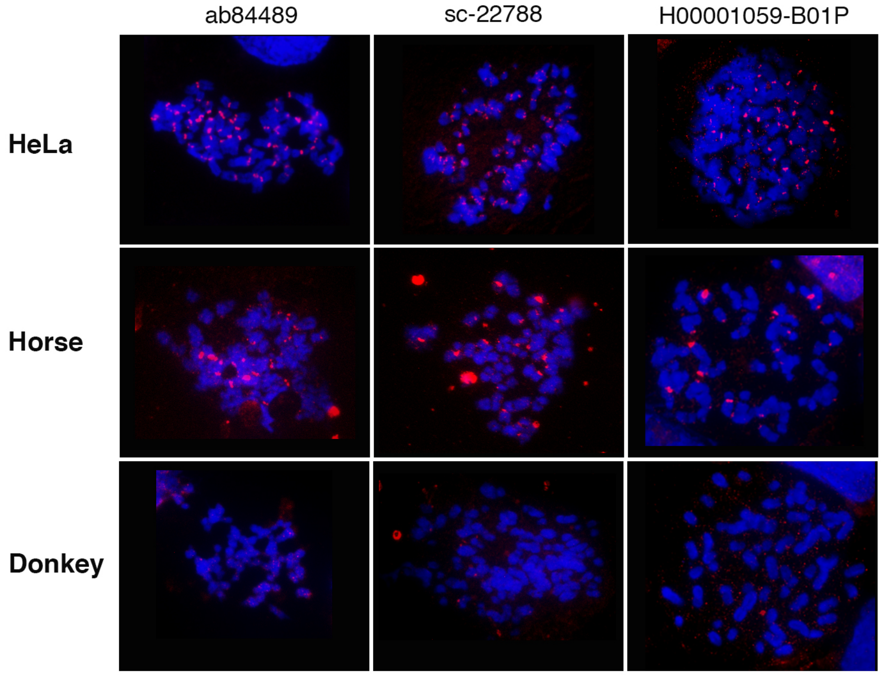


**Figure S5. Immunofluorescence with different anti-CENP-B antibodies on HeLa, horse and donkey metaphase spreads.** The anti-CENP-B antibodies ab84489 (Abcam), sc-22788 (Santa Cruz Biotechnology Inc.) and H00001059-B01P (Abnova) were used.

**Fig. S6**

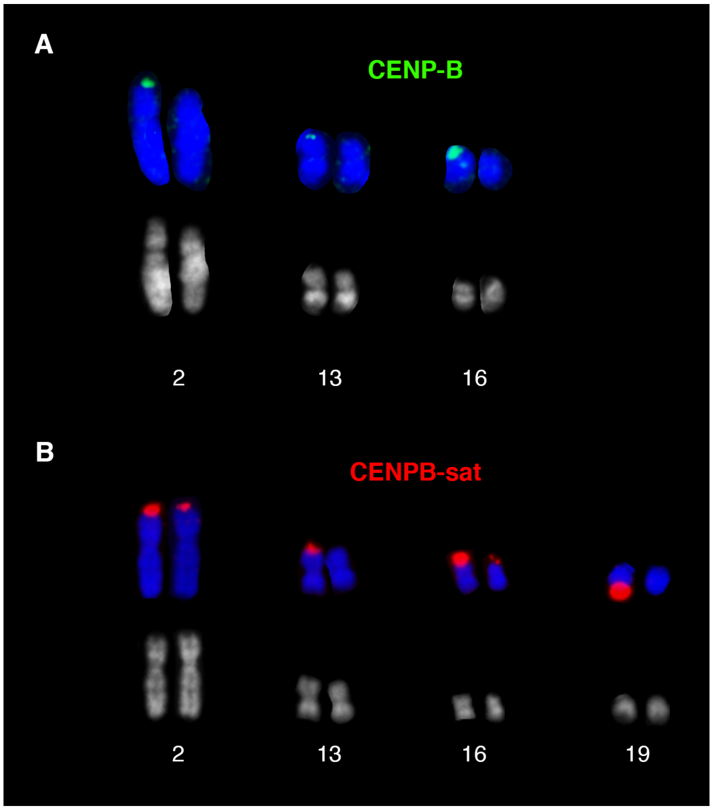


**Figure S6. CENP-B signal polymorphism in Grevy's zebra.** Immunofluorescence with the anti-CENP-B antibody on the homologous chromosome pairs 2, 13 and 16 labelled (A) and fluorescence *in situ* hybridization with the CENPB-sat probe on chromosomes 2, 13, 16 and 19 (B).

**Fig. S7**

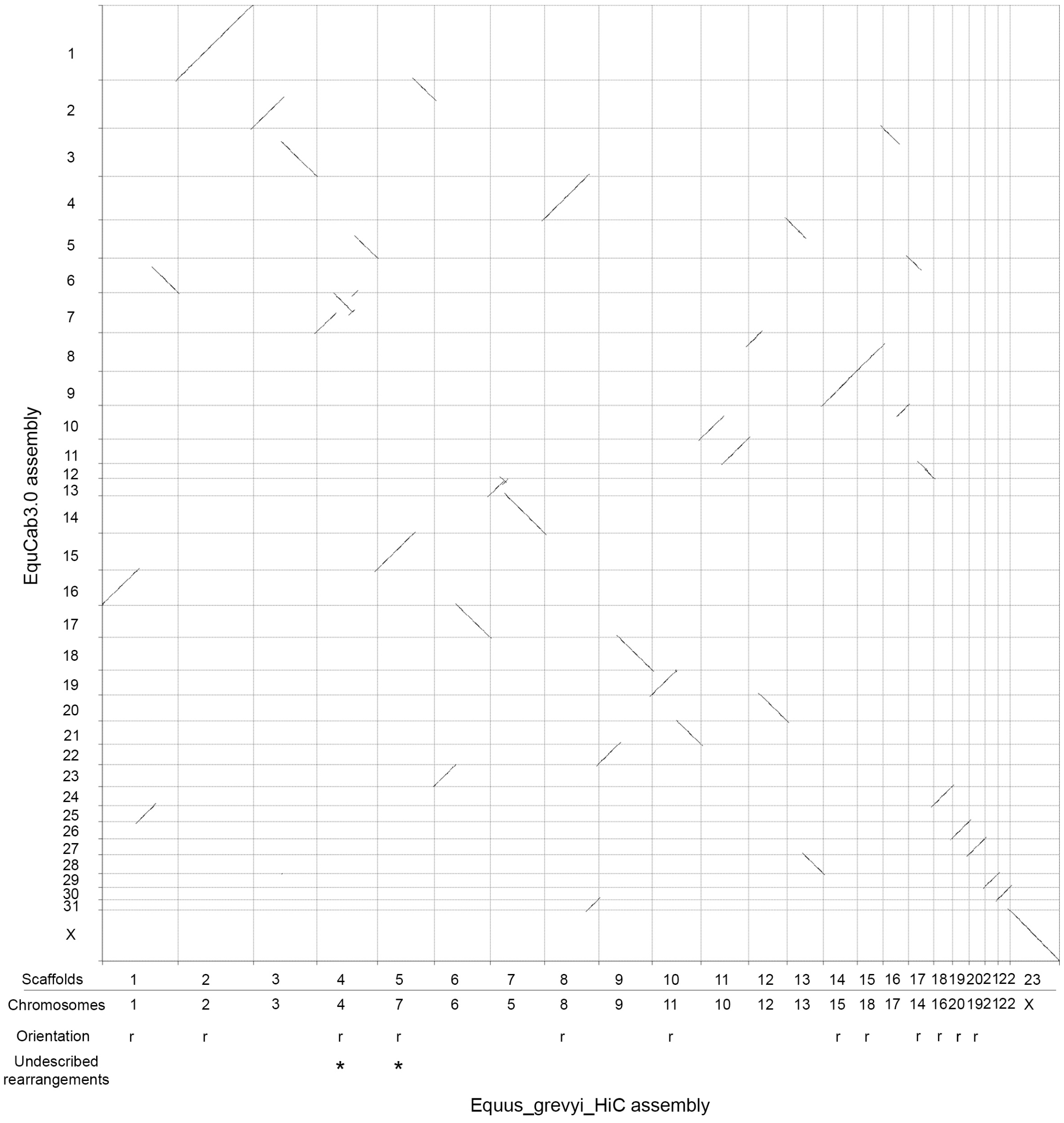


**Figure S7. Pairwise genome comparison between EquCab3.0 and Equus_grevyi_HiC scaffolds.** Aligned segments between horse chromosomes (y-axis) and Equus_grevyi_HiC scaffolds (x-axis) are represented as lines. For each Grevy's zebra scaffold, the chromosome number is reported below. Scaffolds with a reverse orientation compared to the direction previously determined by cytogenetic analysis (Musilova et al. 2013) are indicated with "r". Rearrangements between horse and Grevy's zebra genomes that were not described by Musilova and colleagues are indicated with asterisks.

### Table S1

| Species | Coordinates in genome assemblies and Gene ID | Mutations with respect to the horse (*) |
|---|---|---|
| *E. caballus* | chr22: 19582683-19584500 in EquCab3.0<br>(Gene ID: 100146426) | - |
| *E. asinus* | NC_052191.1 (chr15):19128559-19130376 in ASM1607732v2<br>(Gene ID: 106847668) | Position 150: C>A silent mutation<br>Position 255: C>T silent mutation<br>Position 597: G>C silent mutation<br>Position 824: C>T mutation, alanine-to-valine substitution<br>Position 1288: in-frame GAG insertion (additional Glu residue)<br>Position 1408: in-frame GAG deletion (loss of a Glu residue)<br>Position 1650: C>T silent mutation |
| *E. grevyi* | HiC_scaffold_9: 29914712-29916533 in Equus_grevyi_HiC<br>(https://www.dnazoo.org/assemblies/Equus_grevyi)<br>(Gene ID: not available) | Position 150: C>A silent mutation<br>Position 255: C>T silent mutation<br>Position 597: G>C silent mutation<br>Position 832: C>T silent mutation<br>Position 1288: in-frame GAG insertion (additional Glu residue)<br>Position 1548: G>A silent mutation<br>Position 1650: C>T silent mutation |
| *E. burchelli* | HiC_scaffold_12:77497917-77499777<br>Equus_quagga_HiC<br>(https://www.dnazoo.org/assemblies/Equus_quagga)<br>(Gene ID: not available) | Position 150: C>A silent mutation<br>Position 255: C>T silent mutation<br>Position 597: G>C silent mutation<br>Position 765: C>T silent mutation<br>Position 1288: in-frame GAG insertion (additional Glu residue)<br>Position 1650: C>T silent mutation<br>Position 1713: C>T silent mutation |

**Table S1. *CENP-B* coding sequences.** * The coordinate refers to the nucleotide position of the horse coding sequence.

### Table S3

| **Name** | **Consensus sequence** (the CENP-B box is highlighted in yellow, essential nucleotides for CENP-B binding are written in red) |
|---|---|
| CENPB-sat | TAGGTGCTTTCTGACACTCTCTNNANCCAGTGCACAATGGTGGTTTGTAAAAAGCCTATTGTCTGNC<br>TCCTCCTAAGCATGTGGAAGCACAATCATTTGGGCCTCGNCCCCGTTGCTTAAGGGGAAGATGTAGG<br>CA**TTTCGTCTGAGCCGGGT**TGGCAAGGTGGAAATGTGGCAGAAGCAGAAGTNCCAAAGCTNGNCGAG<br>TTAAGCGCTGAAAAGAAATGGCNTTTCAGCTGCCTTTGTATGAGATGTTCCCAGGNACGCTGTAAGA<br>GCACTGTGGAAAGCGAGTTCTTTCTCAGCTTCCTAAAGAGCTGGANGGCAAGACAGTTTATGGCTTG<br>TCTCCCATTGAAGGATGGAGGCAGTGCTTTGTGCCTTCCACCTCTAGAGCAATGGAGGGCACGGCTG<br>AGAGCAAAGGGGCCTTTCTGACA |

**Table S3. The horse CENPB-sat satellite.**

### Table S4

| | Horse | | | Donkey | | | Grevy's zebra | | | Burchell's zebra | | |
|---|---|---|---|---|---|---|---|---|---|---|---|---|
| | *ChIP* | *Input* | *ChIP/ Input* | *ChIP* | *Input* | *ChIP/ Input* | *ChIP* | *Input* | *ChIP/ Input* | *ChIP* | *Input* | *ChIP/ Input* |
| **CENPB-sat** | 3762.5 | 586.0 | 6.4 | 69.6 | 38.7 | 1.8 | 11756.6 | 1738.2 | 6.8 | 23.6 | 5.7 | 4.1 |
| **201bp-box** | 1509.7 | 196.0 | 7.7 | 14.7 | 7.5 | 2.0 | 3323.0 | 488.5 | 6.8 | 6.9 | 1.4 | 5.0 |
| **ERE-1** | 1154.0 | 1220.6 | 0.9 | 1452.8 | 1298.6 | 1.1 | 1796.3 | 1691.4 | 1.1 | 2182.6 | 1830.3 | 1.2 |

**Table S4. Counts per million (CPM) of ChIP-seq reads obtained with the anti-CENP-B antibody mapped on the CENPB-sat consensus sequence, the 201bp region of CENPB-sat comprising the CENP-B box and ERE-1 retrotransposon.** Enrichment values are reported as ratios between CPM ChIP reads and CPM Input reads.

### Table S5

| | Horse | | | Donkey | | | Grevy's zebra | | | Burchell's zebra | | |
|---|---|---|---|---|---|---|---|---|---|---|---|---|
| | *ChIP* | *Input* | *ChIP/ Input* | *ChIP* | *Input* | *ChIP/ Input* | *ChIP* | *Input* | *ChIP/ Input* | *ChIP* | *Input* | *ChIP/ Input* |
| **CENPB-sat** | 1906.9 | 622.2 | 3.1 | 576.6 | 33.0 | 17.4 | 2874.2 | 1738.2 | 1.7 | 8.1 | 5.7 | 1.4 |
| **37cen** | 151002.7 | 15318.4 | 9.9 | 5921.9 | 2830.2 | 2.1 | 176.7 | 11.0 | 16.0 | 48.6 | 2.9 | 16.9 |
| **ERE-1** | 1216.0 | 1316.7 | 0.9 | 1336.6 | 1275.9 | 1.0 | 1713.7 | 1691.4 | 1.0 | 1883.3 | 1830.3 | 1.0 |

**Table S5. Counts per million (CPM) of ChIP-seq reads obtained with anti-CENP-A antibody mapped on the CENPB-sat consensus sequence, the horse 37cen satellite sequence and ERE-1 retrotransposon.** Enrichment values are reported as ratios between CPM ChIP reads and CPM Input reads.

### Table S8

| | Horse | | | Donkey | | | Grevy's zebra | | | Burchell's zebra | | |
|---|---|---|---|---|---|---|---|---|---|---|---|---|
| **Family** | **Consensus length** | **Read proportion (%)** | **ChIP/ Input** | **Consensus length** | **Read proportion (%)** | **ChIP/ Input** | **Consensus length** | **Read proportion (%)** | **ChIP/ Input** | **Consensus length** | **Read proportion (%)** | **ChIP/ Input** |
| **37cen** | 221 | 1.600 | 9.9 | 221 | 1.80 | 1.5 | 221 | 0.014 | 29.1 | 221 | < 0.001 | 2.9 |
| **2PI** | 44 | 1.100 | 1.4 | 44 | 0.54 | 1.2 | 44 | 0.540 | 1.2 | 220 | 0.110 | 1.1 |
| | 44 | 0.100 | 1.5 | | | | 2037 | 0.017 | 1.0 | | | |
| **CENP B-sat** | 419 | 0.220 | 2.5 | 420 | 0.02 | 14.2 | 419 | 0.660 | 1.8 | 404 | < 0.001 | 2.1 |
| **137sat** | 137 | 0.066 | 1.9 | nd | | | nd | | | nd | | |
| **satA** | 3322 | 0.130 | 1.2 | 4678 | 3.50 | 1.0 | 4681 | 0.630 | 1.6 | 4681 | 0.650 | 1.0 |
| **satB** | 114 | 0.046 | 1.1 | 114 | 0.67 | 1.0 | 114 | 0.390 | 1.0 | 114 | 0.052 | 1.0 |
| **satC** | nd | | | 2467 | 1.00 | 3.3 | nd | | | 4944 | 0.16 | 6.5 |
| **satD** | nd | | | 68 | 0.01 | 0.9 | nd | | | n.d. | | |
| **satE** | nd | | | n.d. | | | nd | | | 75 | 0.025 | 1.0 |

**Table S8. *De novo* identification of satellite repeats in the four species by TAREAN and normalized enrichment values in ChIP-seq reads with anti-CENP-A antibody obtained by ChIP-seq mapper.** nd (not detected)

### Table S10

| Chromosome | Total number of nuclei | Diploid nuclei (%) | Aneuploid nuclei (%) |
|---|---|---|---|
| **Experiment A – normal conditions** | | | |
| ECA9 | 993 | 961 (96.8) | 32 (3.2) |
| ECA10 | 996 | 968 (97.2) | 28 (2.8) |
| **Experiment B – normal conditions** | | | |
| ECA9 | 550 | 539 (98) | 11 (2) |
| ECA10 | 527 | 515 (97.7) | 12 (2.3) |
| **Experiment B – mitotic stress (48 hours 200 nM nocodazole treatment)** | | | |
| ECA9 | 536 | 499 (93.1) | 37 (6.9) |
| ECA10 | 544 | 509 (93.6) | 35 (6.4) |

**Table S10. Interphase aneuploidy analysis of ECA9 (CENP-B negative) and ECA10 (CENP-B positive) chromosomes.**

### Table S11

| Species | Antibody | Read length (bp) | Total number of reads | | SRA accession | |
|---|---|---|---|---|---|---|
| | | | ChIP | Input | ChIP | Input |
| *E. caballus* | anti-CENP-B | 100 | 27,682,380 | 55,359,526 | SRR27295363 | SRR27295364 |
| | anti-CENP-A | 100 | 43,378,388 | 80,130,272 | SRR27325169 | SRR27325168 |
| *E. asinus* | anti-CENP-B | 100 | 29,797,622 | 126,605,282 | SRR27295361 | SRR27295362 |
| | anti-CENP-A | 100 | 44,267364 | 37,434,334 | SRR5515973 | SRR5515972 |
| *E. grevyi* | anti-CENP-B | 125 | 19,020,884 | 24,488,430 | SRR27295360 | SRR17956803 |
| | anti-CENP-A | 125 | 32,468528 | | SRR17956804 | |
| *E. burchelli* | anti-CENP-B | 125 | 26,347,730 | 19,102,434 | SRR27295359 | SRR17956805 |
| | anti-CENP-A | 125 | 24,326822 | | SRR17956806 | |

**Table S11. Antibody, read length, read counts and SRA accession numbers of ChIP-seq experiments**

**Table S12**

| 5'-3' sequence |
|---|
| AAGAATTCGCCACCATGGGCCCCAAGCGGCGGCAGCTGACGTTCC |
| AGGATGGGCCCCAAGCGGCGGCAGCTGACG |
| GTCAAGGGCATCATCCTCAAG |
| TGCTTTCGTGAGGCTGGCTT |
| AAGGATCCTTGCTTTGATGTCCAAGACCCCGAACT |
| CACGCCAGCCGGTCGTACTC |
| GAGGGCAGTGGTGATAGTGG |
| AAGGATCCTTGCTTTGATGTCCAAGACCCCGAACT |

**Table S12. Sequence 5'-3' of primers used in CENP-B CDS amplification and sequencing**
